# Design, Fabrication, and Theoretical Investigation of a Cost-Effective Laser Printing Based Colorimetric Paper Sensor for Non-Invasive Glucose and Ketone Detection

**DOI:** 10.1101/2021.09.23.461386

**Authors:** Manikuntala Mukhopadhyay, Sri Ganesh Subramanian, K. Vijaya Durga, Debashish Sarkar, Sunando DasGupta

**Affiliations:** Department of Chemical Engineering, Indian Institute of Technology Kharagpur, West Bengal – 721302, India; Department of Chemical Engineering, Calcutta University, Kolkata, West Bengal - 700009, India

**Keywords:** Paper device, glucose detection, ketone detection, Richards equation, laser printing, colorimetry

## Abstract

Diabetes, a chronic condition, is one of the prevalent afflictions of the 21^st^ century, and if left unchecked, this ailment could lead to severe life-threatening complications. A widely accepted methodology for monitoring diabetes is the estimation of the glucose and ketone contents in the body-fluids, viz. blood, urine, etc. Additionally, certain conditions such as starvation, and following a protein rich diet (e.g., keto-diet) could also lead to significant changes in the ketone content, thereby resulting in false-positive diagnosis. Hence, a precise, portable, and on-demand procedure for the rapid and combined estimation of glucose and ketone in the bodily-fluids is of utmost importance. To that end, paper-based analytical devices (μPADs) are promising tools, owing to their multitudinous advantages, and compatibility with biofluids. Although, numerous researchers have contributed substantially in the fundamental investigation, design, and fabrication of μPADs for various applications, a combined platform capable of rapid, accurate and on-demand glucose and ketone detection, that is easy to fabricate, is still relatively unexplored. Moreover, the flow dynamics of an analyte, in combination with enzyme-catalysed (for glucose) and uncatalyzed reactions (for ketone), within a porous paper matrix is also vaguely understood. Herein, we present a facile laser-printing based fabrication of colorimetric sensors on a filter paper, for rapid, and non-invasive estimation of glucose and ketone contents in urine. The urine sample, upon being deposited in a particular expanse, is wicked through the paper matrix, and reacts with specific reagents in the designated zone(s), giving rise to a final color, concomitant with the glucose or ketone content in the sample. The device design enables the liquid to be wicked into the porous matrix in a way that would concentrate the colored product in a dedicated detection zone, thereby augmenting the feasibility for accurate colorimetric detection. Furthermore, we present for the first time, a detailed dynamic model of the flow-field in a variable cross-section paper device using the Richards’ equation, while also considering the species transport and reaction kinetics within the porous media. The results of the numerical simulation agree well with those observed experimentally, thereby validating the present model. Finally, we also developed a web and desktop-based application that would enable the user to upload the images of the colored zones to provide an accurate estimate of the glucose and ketone content in the sample. We believe that our model, in combination with the proposed fabrication methodology, and the in-house developed app., would enable rapid and reliable fabrication of μPADs for various fundamental investigations, and applications pertaining to affordable health-care monitoring.

**Graphical Abstract:** 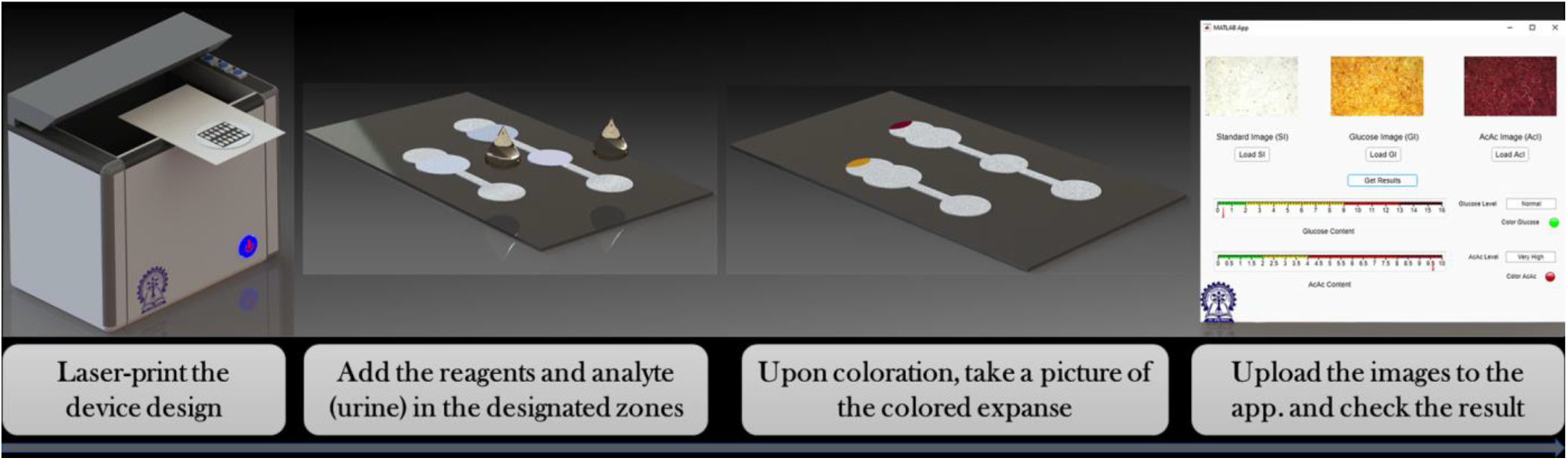

## 1. Introduction

The examination and analysis of various body fluids such as blood, urine, saliva, etc., has played a vital role in medical diagnosis of various ailments, for over several centuries. (Chen et al., 2016; Hu et al., 2014; Parsa et al., 2018; Sefiane, 2010) The concentration of diverse chemical and biochemical species in the body fluids viz. proteins, drugs, glucose, inorganic ions, ketones, lactate, uric acid etc., can serve as potential biomarkers in determining the health status of a subject. (Gabriel et al., 2017; Klasner et al., 2010; Lim et al., 2017; C. C. Lin et al., 2011; Soni and Jha, 2015) With advancements in global health becoming imperative for economic growth especially in developing nations, the emergence of microfluidic platforms have demonstrated significant promise towards the development of affordable diagnostics (Ballerini et al., 2012; Brutin et al., 2011; Campbell et al., 2018; Cate et al., 2015; Chin et al., 2007; Li et al., 2012; Mao and Huang, 2012; Martinez et al., 2010b; Mukhopadhyay et al., 2020; Yetisen et al., 2013). One such paradigm is the evolution and proliferation of paper-based analytical devices (μPADs). These multifunctional systems have piqued a lot of interest amongst researchers owing to their several desirable properties such as cost-effectiveness, ease of availability, chemical inertness, compatibility with organic, inorganic, and biological entities, and biodegradability, to name a few. (Chen et al., 2015; Dou et al., 2015; Kumar et al., 2019; Martinez et al., 2007) In the preceding decade, paper has been immensely used as a substrate for multitudinous applications such as colorimetric detection (Gabriel et al., 2017; Klasner et al., 2010; Shen et al., 2012), electrochemical and biological sensing (Ahmed et al., 2016; Nie et al., 2010), bioanalysis such as estimation of glucose, protein, and urea content in urine (Lepowsky et al., 2017; Lim et al., 2017; Martinez et al., 2008a; Tai et al., 2021), identifying biomarkers for disease detection (Campbell et al., 2018; Fu and Wang, 2018; Islam et al., 2018; Tai et al., 2021), personalised health care (Mahato et al., 2017; Yetisen et al., 2013), Point-of-Care (PoC) medical diagnosis (Akyazi et al., 2018; Cate et al., 2015; Gong and Sinton, 2017; Hu et al., 2014; Li et al., 2012; Yamada et al., 2017) and even for crime scene detections (Musile et al., 2021). One of the versatilities of a paper-microfluidic device is the possibility to combine the platform with several detection strategies, such as electrical impedance, conductance, spectrometry, etc. (Abe et al., 2008; Ahmed et al., 2016; Kap et al., 2021; Nie et al., 2010; Selvakumar and Kathiravan, 2021; Zhang et al., 2021) Colorimetric detection coupled with μPADs is also one of the most favoured methods for development of several PoC devices, such as monitoring the nitrate content in human saliva (Bhakta et al., 2014), analysing tear electrolytes (Gabriel et al., 2017; Kang et al., 2017; Yetisen et al., 2017), detection of clinically relevant analytes in urine and other biofluids (Gabriel et al., 2017; Islam et al., 2018; Klasner et al., 2010; Park et al., 2005; Sechi et al., 2013; Soni and Jha, 2015; Zhang et al., 2021) etc., due to its cost-effectiveness, portability, and user-friendliness. In this regard, a significant quantum of research efforts was directed towards simple and sustainable fabrication of paper microfluidic devices using traditional and novel techniques such as wax, inkjet, and flexographic printing (Kumar et al., 2019; Nishat et al., 2021; Olkkonen et al., 2010; Xia et al., 2016), plasma treatment (Li et al., 2008), photolithography (Whitesides, 2006), wet etching (Cai et al., 2014), vapour phase deposition (Demirel and Babur, 2014), and laser treatment (Yetisen et al., 2017). Although most of the abovementioned methods are fast and straightforward, they either require expensive equipment(s), thereby, leading to an increase in the overall cost of the device; or necessitate the usage of toxic chemicals, thus, rendering the final device incompatible with organic solvents (Kumar et al., 2019). Hence, a modified fabrication methodology, which is simple, cost-effective, and does not require the usage of any toxic chemicals is of utmost importance, for continued fundamental exploration, and large-scale adaptation of paper-based microfluidic devices.

One of the prevalent maladies plaguing the 21^st^ century is diabetes and therefore, close monitoring of glucose and ketone levels, in conjunction with certain lifestyle modifications is extremely crucial for diabetic patients, failing which could result in severe complications such as diabetic ketoacidosis (DKA), stroke, blindness, kidney failure, and limb amputations (WHO Global Report on Diabetes, 2016). DKA is pre-eminent in type 1 diabetic patients and if left untreated can often lead to adult respiratory distress syndrome, cerebral edema, acute kidney failure, coma, blindness, fatigue, and in some serious cases, could even cause death. (Brink, 1999; Dunger et al., 2004; Foster et al., 2011; Kerl, 2001; Wallace and Matthews, 2004; Wolfsdorf et al., 2006) Additionally, the concentration of ketone bodies (β-hydroxy butyrate (BHB) and acetoacetate (AcAc)) may also increase, because of starvation and consumption of low carbohydrate diets (viz. keto-diet), etc.), (Laffel, 1999) thus, making it crucial to monitor both the ketone and glucose contents simultaneously to prevent any false positive diagnosis of diabetes. At present, one of the widely used and cost-effective devices for self-monitoring of glucose and ketone contents are the dipsticks (Sharp et al., 2008) which often produces erroneous results due to inaccurate eye estimations. Additionally, the detection range of the glucosticks starts from 5.6 mM (100 mg/dL) of glucose, whereas the clinical range of glucose indicating abnormality begins from 2mM (36 mg/dL) (Martinez et al., 2007, 2008a; Sechi et al., 2013; Sharp et al., 2008). Therefore, in the recent years, there is a significant surge of research articles describing various detection strategies using paper-based devices for successful estimation of glucose content in biofluids.(Aksorn and Teepoo, 2020; Cao et al., 2020; Chaiyo et al., 2018; Chauhan and Toley, 2021; Chen et al., 2012; Gu et al., 2019; Kap et al., 2021; Kim et al., 2020; Lamas-Ardisana et al., 2018; Rossini et al., 2019) However, as stated previously, a thorough estimation of the ketone content is also critical for proper healthcare monitoring in diabetic patients,(Khan et al., 2004; Klocker et al., 2013; Laffel, 1999; Verma and Sharma, 2010) but the research pertaining to detection of ketone bodies in body fluids, using μPADs is still nascent.(Klasner et al., 2010; Martinez et al., 2010a)

The above studies clearly underscore the remarkable advancements that have been made in the preceding decade towards the development of paper-based analytical devices. (Martinez et al., 2010a; Nishat et al., 2021; Selvakumar and Kathiravan, 2021; Svendsen, 2015; Yetisen et al., 2013) However, the fabrication of a facile, low-cost, and non-invasive paper-based platform, capable of simultaneously detecting the glucose and ketone contents, still remains relatively unexplored. To that end, in the present work, we have designed and created a paper device on a Whatman® filter paper with two unique channels for simultaneous self-monitoring of glucose and ketone (AcAc) content in urine, which are significant biomarkers for the diagnosis of DKA (Khan et al., 2004; Klasner et al., 2010; Klocker et al., 2013; Laffel, 1999; C. Lin et al., 2011; Martinez et al., 2010a; Molina et al., 1989). The fabrication of the hydrophobic barriers on the paper device was accomplished by the systematic deposition of printing ink using a commercial laser printer. The urine sample, wicks through the porous matrix, and reacts with the reagents in a specific expanse to produce a distinct color in a designated zone, concomitant with the content of the glucose or ketone in the sample. Additionally, it is also well overt that, a significant quantum of research efforts has been dedicated to the mathematical understanding of the flow through a porous media in general, and a paper matrix in particular.(Buser, 2016; Cai and Yu, 2011; Chang et al., 2018; Chauhan and Toley, 2021; Elizalde et al., 2015; Modha et al., 2021; Rath et al., 2018; Rath and Toley, 2021; Starov et al., 2002) However, there is a paucity in literature pertaining to the theoretical comprehension of the flow-assisted reaction dynamics and the associated species transport through a paper-based sensor with variable cross-section along the flow path. Herein, we attempt to bridge the gap by not only carefully designating and fabricating the paper devices using a commercially available laser printer, but by also performing a systematic study of the hydrodynamics within the paper matrix by using the Richards’ equation.(Richards, 1931) Furthermore, we present for the first time, the flow modulated coloration of the final product in the enzyme catalyzed oxidation of glucose, and the non-catalyzed reaction of the ketone molecules, in porous filter paper devices with variable cross-section, within the framework of a finite element model using COMSOL Multiphysics®. Our theoretical results clearly demonstrate the dominant mechanisms involved in the formation of the colored products in both the devices at the respective zones, as observed experimentally, thereby highlighting the effectiveness of the numerical simulations. Additionally, we have also developed an in-house web and desktop application using MATLAB® which would enable the user to take a picture of the colored expanse in the two sensors, and upload it to the app., thereby allowing rapid and on-demand diagnosis. We demonstrate that our paper-based devices and the associated numerical simulations, and the in-house developed app., could be used to develop a potential prototype of a Point-of-Care device capable of detecting both glucose and ketone (AcAc), with adequate accuracy and minimal training. We believe that the present study could aid in minimizing the time and effort required in design and fabrication of a paper-based sensor, thereby finding application in low-resource settings, for continuous, reliable, and affordable health-care monitoring.

## 2. Materials and Methods

### 2.1. Materials and Reagents

All the chemicals used in the present study were purchased from Sisco Research Laboratories Pvt. Ltd. (SRL), India, unless mentioned otherwise. All the filter papers used were manufactured by Whatman®.

### 2.2. Preparation of Artificial Urine

With elevated levels of glucose and ketone concentrations, the body tries to maintain homeostatis, which is a steady state of internal physical and chemical conditions, thus, disposing the excess glucose and ketone contents via urine. Moreover, ketones appear in a subject’s urine prior to their detection in blood (Klasner et al., 2010; Sharp et al., 2008). Hence, regular monitoring of glucose and ketone contents in the urine is an efficient, non-invasive, and cost-effective way for early detection of diseases (Martinez et al., 2008a, 2010b).

To standardize the present methodology, artificial urine was considered as a model system to ensure repeatability and to establish a standardization curve, which could potentially be used to analyse actual diseased samples. The artificial urine was prepared following the standard recipe as described by Brooks and Keevil (Brooks and Keevil, 1997; Zhang et al., 2019), wherein 1.1 mM lactic acid, 2.0 mM citric acid, 2*5* mM sodium bicarbonate, 170 mM urea, 2.*5* mM calcium chloride, 90 mM sodium chloride, 2 mM magnesium sulfate, 10 mM sodium sulfate, 7 mM potassium dihydrogen phosphate, 7 mM di-potassium hydrogen phosphate, 2*5* mM ammonia chloride, 1 M hydrochloric acid were dissolved in 1 L of distilled water. All the ingredients were stirred using a magnetic stirrer for 10-1*5* minutes until all the contents were completely dissolved in the solvent. The artificial urine was then stored at 2 °C and all the experiments were performed within a week of its preparation. However, regular checks for depositions were done to monitor the stability of the same.

### 2.3. Glucose Assay

The glucose assay was performed following the established procedure as reported in the literature.(Klasner et al., 2010; Martinez et al., 2008a, 2008b, 2007) When glucose reacts with glucose oxidase in the presence of oxygen, it forms glucono delta-lactone and hydrogen peroxide as depicted in equation 1. The produced hydrogen peroxide further reacts with potassium iodide in the presence of horseradish peroxidase with simultaneous oxidation of iodide to iodine (equation 2), resulting in golden-brown coloration. The intensity of the color was then quantified and correlated with the concentration of the glucose. The glucose content in urine above ~2 *mM* is typically an indicator of diseases (Martinez et al., 2008a, 2007; Sechi et al., 2013), and hence the glucose content in the present study was varied between 1 to 1*5* mM to accurately establish a calibration curve, to detect and delineate samples that are healthy and otherwise.

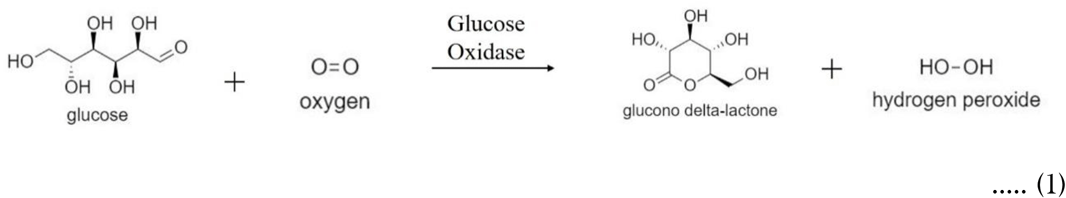

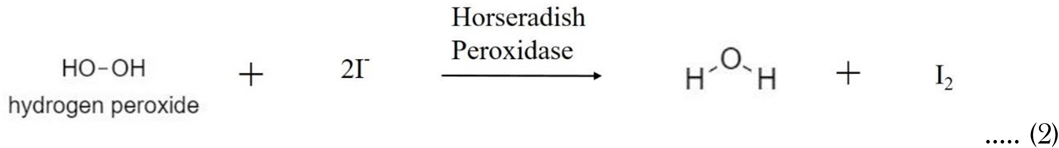

### 2.4. Ketone Assay

The acid form of acetoacetate, which has similar properties to the ketone form, was synthesized by hydrolysing its ester form, i.e., ethyl acetoacetate (Clayden et al., 2001; Hawk, 1955; Pedersen, 1961), as shown in equation 3. The standard sodium nitroprusside test was followed to detect the acetoacetic acid content in the urine. (Hawk, 1955; Klasner et al., 2010) AcAc reacts with glycine to form an imine derivative (equation 4) which further reacts with sodium nitroprusside to produce a magenta colored complex (equation *5*). The intensity of the magenta color was then quantified and correlated to the concentration of the ketone present in the urine sample. Similar to glucose, the presence of AcAc at concentrations greater than ~2 *mM* in urine, could be deemed to be an indicator of unhealthy levels of ketone moieties in the body (JP and AJ., 1990; Sharp et al., 2008). Hence the concentration of AcAc was varied from 1 to 10 mM in the present study, to accurately delineate different bands corresponding to the ketone content in urine.

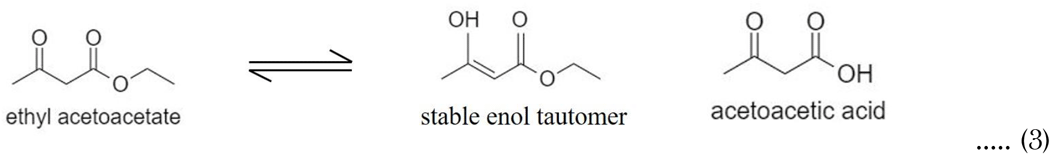

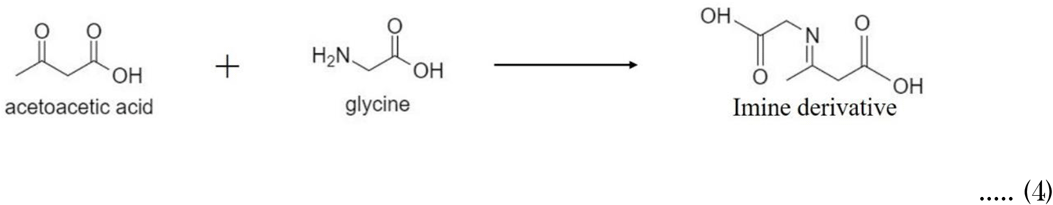

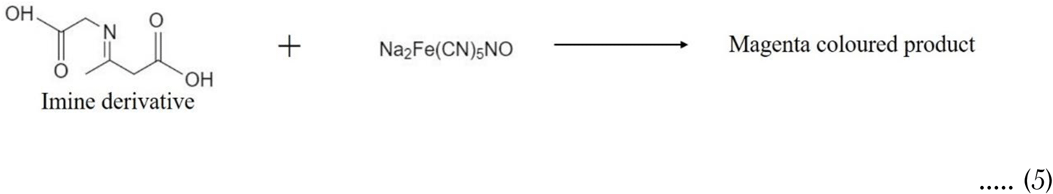

## 3. Experimental Procedures

### 3.1. Optimizing the Substrate

Five different grades of Whatman filter paper - Grade 113, Grade 43, Grade 4, Grade 3, and Grade 1 were initially selected as possible substrates for the present work (the detailed characteristics of all the chosen substrates are presented in S1 of the ESI). The time required for the complete formation of the hydrophobic barriers coupled with the ease of printing and the overall cost of the prepared substrates played a significant role in selecting the most appropriate grade for the process. Amongst all the chosen filter papers, Grade 1 was finalised as the choice of substrate for the present work due to its ease of formation of hydrophobic barriers within the least amount of time (circa. 30 minutes).

### 3.2. Fabrication of the Devices

The channels were created on the paper substrate utilizing a commercially available desktop printer (HP laser Pro 400 series) without any specific customizations. Initially, an array of the designs was created to print several devices simultaneously, to ensure maximum utilization of the substrate surface. A mirror image of the array was then generated and using the laser printer, the original array was printed on one side of an A4 sheet paper, while the mirror image was printed on the other side to perfectly align the designs on both sides. It is extremely crucial to get perfect alignment on both sides, to avoid uneven melting of the ink and improper formation of hydrophobic barriers, thereby, resulting in leakage of the fluid through the channels. Upon perfect alignment, the printed portion of the array was peeled off, and a Whatman grade 1 filter paper was pasted on the cut-out part. Next, the printing process was repeated on the filter paper, which was then removed from the A4 sheet and heated on a hot plate at 180°C for about 40 minutes. A copper block of thickness 1.*5* cm, preheated to 180°C was placed on top of the substrate so as to heat it from both sides, thus, allowing uniform melting of the ink. The ink, mostly consisting of carbon black, further percolated into the microfibres of the paper, filling the entire thickness of the substrate leading to successful creation of the channels on the filter paper. Upon completion of the heating, the devices were carefully removed from the hot-plate and placed in a vacuum desiccator until further use to ensure negligible contamination. Before depositing the analyte, precise quantities of specific reagents pertaining to the glucose and ketone assay, as stated previously, were carefully deposited in the designated zones, and the solvent (water) was allowed to evaporate. The final device with the prescribed quantity of reagents, could now be used for the accurate estimation of glucose and ketone content in a given urine sample. A pictorial representation of the steps followed for fabricating the paper-based device is depicted in Figure 1.

**Figure 1:**
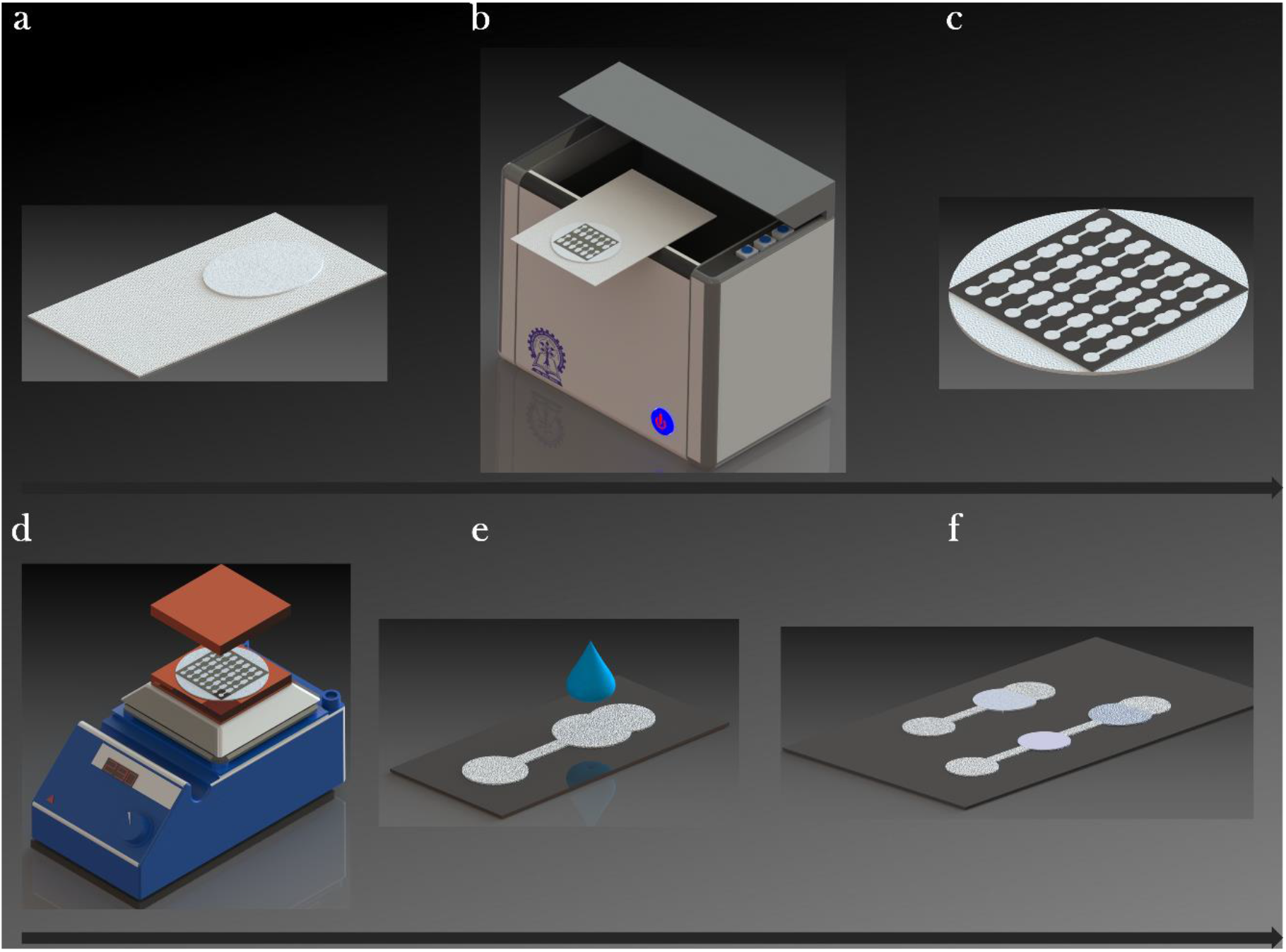
Schematic depicting the detailed procedure followed for laser-printing the device designs onto a Whatman grade-1 filter paper. From a through f, the filter paper was carefully placed atop an A4-size sheet, and the premade designs were printed on to the filter paper following symmetry and proper alignment protocol. The substrates were then assiduously peeled-off from the A4 sheet, and sandwiched between a hot-plate and a copper block, both pre-heated to 180 °C, to ensure thorough melting of the printer-ink. After heating the substrate for circa. 40 minutes, the devices were carefully removed from the heater and allowed to cool down. The devices were stored in an inert ambiance until use, to ensure negligible contamination from dust and other foreign objects. Subsequently, specific reagents pertaining to glucose and ketone assays were deposited in the designated zones of the respective devices, and were allowed to dry. The final devices were used for the successful estimation of glucose and ketone contents in the urine samples.

### 3.3. Device Design Optimization and Characterization for Glucose Assay

It is well established that a value exceeding 2mM (approx.) of glucose content is urine is a widely accepted indicator of diseases (Martinez et al., 2008a, 2007; Sechi et al., 2013). However, the commercially available glucose detection strips can detect the same only if the content is above ~6 *mM* (Sharp et al., 2008). Moreover, inaccurate eye-estimations make these methods quite unreliable and erroneous. Therefore, our target was to develop a device which could detect the cut-off glucose concentration (~2 *mM*), so that these devices could be used for rapid initial screening with adequate accuracy.

A wide array of designs was initially considered for the glucose assay, where each new iteration was a result of modifications to the previous one based on the observations (detailed modifications for each device generation are presented in S2 of the ESI). The final design that was chosen for performing the glucose assay is depicted in Figure 2 (a and b).

**Figure 2:**
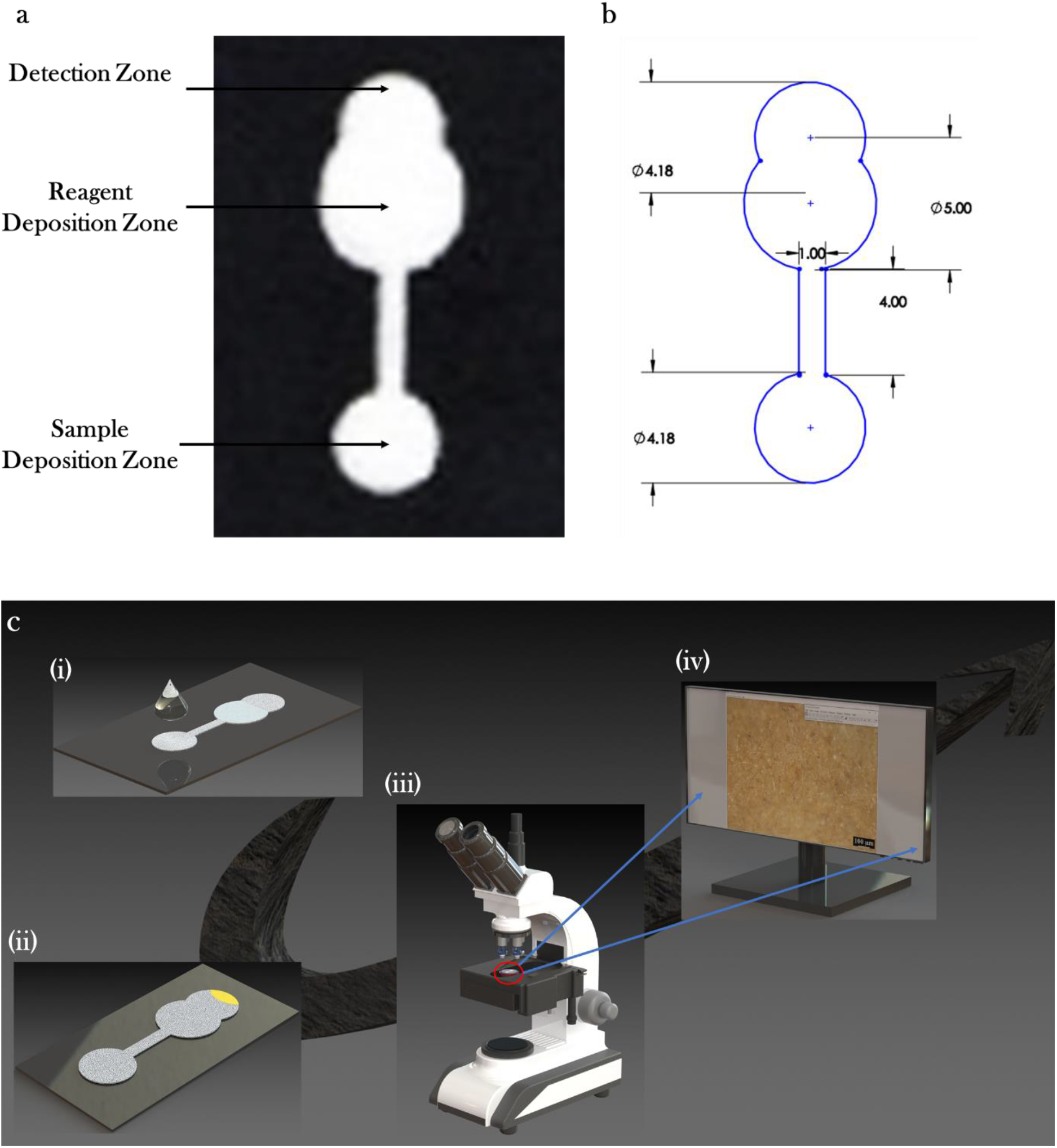
(a) Pictorial representation of the final channel design that was chosen for performing the glucose assay. (b) Dimensional representation of the channel design. (All dimensions are in mm). (c) Schematic representation of the procedure adopted for performing the glucose assay. From (i) through (iv), the glucose sensor with reagents in the designated zone was carefully removed from the vacuum desiccator, and a *5* μL droplet of the analyte (urine) was deposited in the sample zone. The urine wicked through the paper matrix, and travelled along the flow-path and reacted with the reagents in the respective zone, resulting in the formation of the colored product, which was carried along the device to the detection zone. After 30 min. (approx.), the paper-based sensor was placed on the stage of an upright microscope and a high-resolution image of the colored expanse was captured for further analysis. The image was uploaded into an open-source software (ImageJ), and the resultant intensity (gray-values) was captured for each sample with glucose concentration varying from 1 mM to 1*5* mM. The data obtained thereafter was further processed, resulting in the final standardization curve.

For sample preparation, dextrose was added to the artificial urine in varied concentrations from 1 mM to 1*5* mM. All samples were stored at 2 °C and were used within 3 days of preparation. The first reagent consisted of 0.3 M trehalose and 0.6 M potassium iodide dissolved in 100 mM phosphate buffer at pH 6.0. (Klasner et al., 2010) For the second reagent, one part of 30 unit/mL horseradish peroxidase was mixed thoroughly with five parts of 120 units/mL glucose oxidase (Klasner et al., 2010). To perform the assay on the prepared channel, a *5* μL droplet of the first reagent was spotted on the reagent deposition zone and was allowed to dry. Following this, another *5* μL drop containing the second reagent was placed on top of reagent 1 and was left for drying.

Upon requirement, a *5* μL droplet of the analyte was spotted on the sample zone to ensure that the urine wicked through the channel and reacted with the reagents. The detection zone was then observed (after ~30 minutes) under an upright microscope (LEICA DMLM6000M) in reflection mode using a 10x objective lens and high-quality images were captured, as depicted in Figure 2(c). The golden-brown color observed in the detection zone was consistent almost for the next two hours. These images were then analysed using an open-source software viz. ImageJ (Fiji, v. 2.0) to quantify the intensity of the developed color. This methodology was repeated for all concentrations of glucose (1 mM to 1*5* mM) and for each concentration, the process was repeated at least 30 times to correlate the intensity of the developed color with the glucose content in urine. All experiments pertaining to the two assays were performed under ambient conditions where the room temperature was maintained at 27 ± 3 °C and the relative humidity was ~ 40–55 %.

Though the use of cell phone cameras would appear to be a convenient medium in the development of several point of care devices, we preferred capturing the images using an upright microscope for the standardisation process, as often the images clicked by the cell phone cameras are less reliable and are of lower resolution (Kaewarsa et al., 2017; Soni and Jha, 2015). Moreover, each phone has disparate cameras based on its brand and price-range, thus, rendering them unsuitable for the standardisation process. Additionally, most widely used camera phones cannot focus on objects closer than ~ 20 cm and are unable to provide high quality images of small test zones. (Gabriel et al., 2017; Martinez et al., 2008a) Thus, a systematic investigation is essential to make these systems reliable enough for analysis in point-of-care devices. It must be highlighted that the current aim of the work is to develop a cost-effective method which can be instrumental in the design of a non-invasive point-of-care device that can accurately detect both higher contents of glucose and ketone and can assist in the initial screening of the un-healthy samples, which, if required, can be further followed by costlier and labour-intensive tests. However, to facilitate the applicability of our study to different visualization platforms (viz. scanner or other cameras) and to make the results independent of a microscope, we have also developed a web and desktop application, the details of which are presented subsequently.

### 3.4. Device Design Optimization and Characterization for Ketone Assay

Unlike the glucose assay, the detection of ketone in urine (Hawk, 1955) required two distinct zones so that the target molecule (acetoacetate) could wick through the channel, react with glycine in the first reagent zone and form an imine derivative. The imine derivative on further wicking would react with sodium nitroprusside in the second reagent zone and produce a magenta-colored product in the detection zone. The presence of two zones is essential as no color change would be perceptible if either acetoacetate or glycine reacts with sodium nitroprusside. (Klasner et al., 2010) Therefore, it was imperative that the two species reacted prior to coming in contact with sodium nitroprusside to develop the magenta color. The channel design was kept similar to that already optimized for the glucose assay, only with the addition of another reagent zone, as depicted in Figure 3 (a and b).

**Figure 3:**
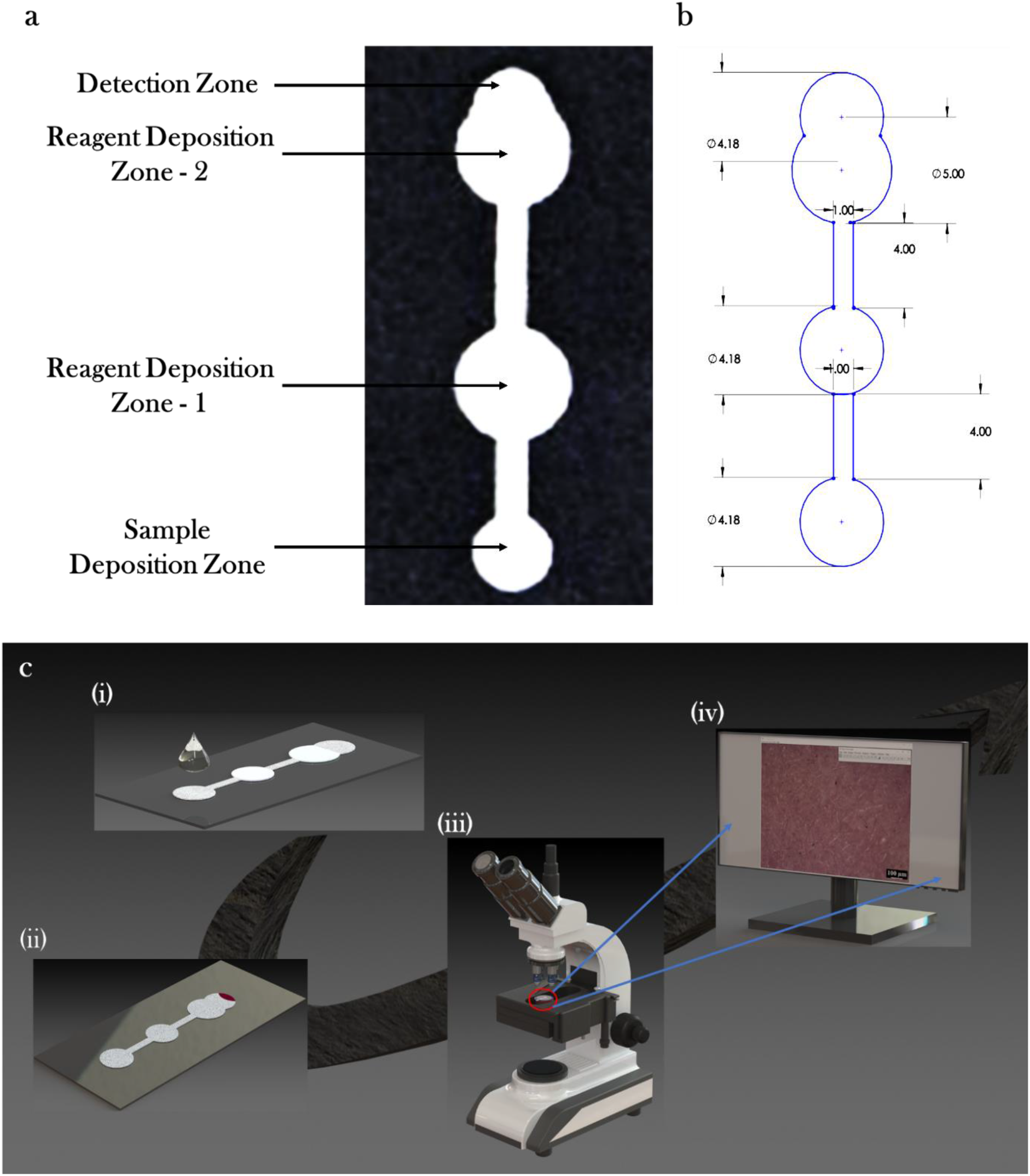
(a) Pictorial representation of the final channel design that was chosen for performing the AcAc assay. Since the presence of two zones was vital for the formation of the colored product, the design for the ketone sensor was slightly modified by the addition of another reagent zone, which would enable the wicking of the product from the first zone to the second. (b) Dimensional representation of the channel design. (All dimensions are in mm). (c) From (i) through (iv), the ketone sensor with reagents in the designated zone was carefully removed from the vacuum desiccator, and a 10 μL droplet of the analyte (urine) was deposited in the sample zone. The urine wicked through the paper matrix, and travelled along the flow-path and reacted with the reagents in the two respective zones, resulting in the formation of the colored product, which was carried along the device to the detection zone. After 30 min. (approx.), the paper-based sensor was placed on the stage of an upright microscope and a high-resolution image of the colored expanse was captured for further analysis. The image was uploaded into an open-source software (ImageJ), and the resultant intensity (gray-values) was captured for each sample with AcAc concentration varying from 1 mM to 10 mM. The data obtained thereafter was further processed, resulting in the final standardization curve.

For sample preparation, AcAc was added to the artificial urine samples giving rise to a resultant ketone concentration ranging from 1 mM to 10 mM. All samples were stored at 2 °C and were used within 3 days of preparation. Reagent 1 consisted of 100 mM glycine in 100 mM phosphate buffer at a pH of 9.4. The second reagent was *5*% dimethyl formamide (DMF) (v/v) and *5*% sodium nitroprusside (w/w) in water (Klasner et al., 2010).

On the prepared channel, a *5* μL droplet of reagent 1 and reagent 2 were deposited on the first and second zones respectively and were allowed to dry. Upon requirement, 10 μL of the analyte (urine) was spotted on the sample zone for the urine to wick through the channel and react with the reagents simultaneously. The detection zone was then observed (after ~ 30 minutes) under an upright microscope (LEICA DMLM6000M) in reflection mode using a 10x objective lens and high-quality images were captured, as depicted in Figure 3(c). The magenta color observed in the detection zone was consistent almost for the next two hours. The procedure that was followed for capturing and analysing the images (for all the concentrations from 1 mM to 10 mM) was same as that of the glucose assay (outlined in the previous sub-section). Figure 3 (c) depicts the entire procedure that was being employed for performing the ketone assay.

It must be highlighted here that the detection of glucose and acetoacetate are two mutually exclusive processes. The reagents chosen herein were to ensure that the presence of glucose or any other analyte in urine would not affect the results of the ketone assay because, the other analytes are unreactive to the chemicals required for the assay and do not produce any other colored products. The same rationale is applicable for the reagents that have been used for AcAc detection as well. In this work, as stated previously, artificial urine has been used which has the same composition as natural urine and none of the other components present in the urine have been found to affect the assays. Hence, we affirm that the results of the present work could be extended to detect the glucose and ketone content in actual urine samples, without loss of generality.

## 4. Theoretical Framework

The following section provides a detailed outline of the system description, the assumptions, and the governing equations used to solve the fluid flow mediated transport of different species, and their reaction kinetics within a porous filter paper.

### 4.1. Fluid Flow Through the Paper Matrix

The flow of a fluid through a porous media has long intrigued researchers, owing to the complex interplay of various pertinent forces, which are non-prevalent on a non-porous substrate. (Buser, 2016; Cai and Yu, 2011; Chang et al., 2018; Chaudhury et al., 2016; Elizalde et al., 2015; Kar et al., 2020; Liu et al., 2015; Millington, 1959; Millington and Quirk, 1959; Modha et al., 2021; Richards, 1931; Shou et al., 2014) The pioneering work by Lucas (Lucas, 1918) and Washburn (Washburn, 1921) over a century ago, which consisted of modelling the porous media as a bundle of cylindrical capillary tubes, is still widely used today, even for the description of flow through a paper matrix, owing to its simplistic relationship between the time-scale and the liquid flow-front. (Buser, 2016; Cummins et al., 2017; Kar et al., 2020; MacDonald, 2018; Modha et al., 2021; Patari and Mahapatra, 2020) However, such a consideration suffers from significant limitation, especially while dealing with the intricate and tortuous nature of the fibres in a paper media. (Buser, 2016; Cummins et al., 2017; MacDonald, 2018; Modha et al., 2021) Therefore, as an alternative, researchers used the Darcy’s law for modelling the fluid imbibition in a paper device. (Buser, 2016; MacDonald, 2018; Modha et al., 2021) Although this approach could be used to model two-dimensional porous systems without any loss of generality, certain situations involving unsaturated flow in a paper matrix, could not be efficiently modelled by this approach. (Buser, 2016; MacDonald, 2018; Modha et al., 2021) Hence, in the recent years, researchers have taken cues from hydrology, and have gradually segued towards the use of Richards’ equation in the modelling of partially saturated fluid flow in porous media. (Buser, 2016; Chauhan and Toley, 2021; Rath et al., 2018; Rath and Toley, 2021) This equation, which was proposed by Richards in 1931, (Richards, 1931) is a modified form of the Darcy’s equation, and relates the changes in the hydraulic properties of the porous medium with the fluid flow as the liquid fills some pores and drains the others. (Buser, 2016; Modha et al., 2021; Rath et al., 2018; Rath and Toley, 2021) Such modelling approaches are critical in designing paper-based devices, owing to the time-varying nature of the fluid imbibition, which in turn impacts the chemical kinetics of reacting species, the device dimensions, and the analyte residence time within the porous media, to name a few. (Buser, 2016; Chauhan and Toley, 2021; Rath and Toley, 2021) Hence, a sufficient understanding of the physics governing the fluid flow in a μPAD, would aid in designing efficient sensors capable of numerous functionalities. Herein, we use the Richards’ equation for modelling the fluid imbibition in the paper matrix, which is represented as follows (Rath et al., 2018; Rath and Toley, 2021):

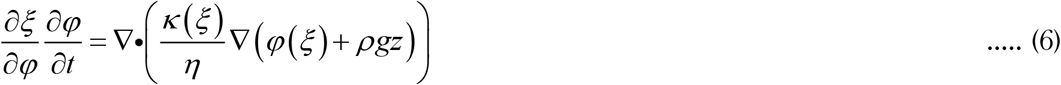

where, ξ denotes saturation, which is defined as the fraction of the total volume of the porous paper matrix, occupied by the fluid (analyte) [0 ≤ ξ ≤ *ε*], *ε* is the porosity of the filter paper, *φ* denotes the capillary pressure, *t* represents the time, *η* is the liquid viscosity, and *κ* is the permeability. Assuming that, *z* denotes the direction normal to the surface of the porous paper, we can safely neglect the *ρgz* term in the above equation, because the two sensors were placed horizontally atop a rigid surface throughout the length of the experimentation.

Given the non-linear relationship exhibited by *κ*(ξ) and *φ*(ξ) in the above equation, several correlations have been proposed to obtain a physically relevant solution to the Richards’ equation. (Buser, 2016; Modha et al., 2021; Rath et al., 2018; Rath and Toley, 2021) One such widely accepted formulation is that proposed by van Genuchten, (Buser, 2016; Rath et al., 2018) and is represented as follows (Rath et al., 2018):

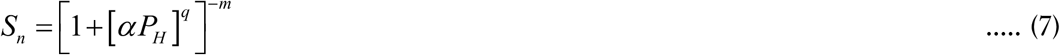

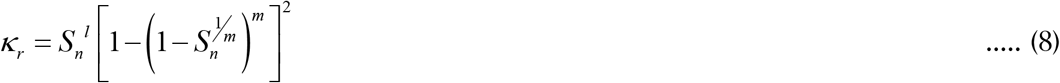

where, *α*, *q*, and *l* denote the van Genutchen parameters, and for the Whatman grade-1 filter paper used in the present work, these values are available in the literature (Buser, 2016; Chauhan and Toley, 2021; Rath et al., 2018). 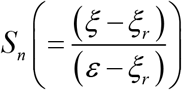 denotes the normalized saturation, 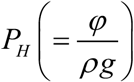 represents the pressure head, 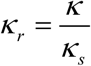, *K_s_* is the value of permeability at complete saturation (i.e. *S_n_* = 1), and 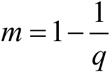.

To model the physics of fluid flow within the porous media, we used COMSOL Multiphysics (v. *5*.4). The Richards’ equation is a versatile interface within the “Subsurface Flow module”, and the governing equation (equation (6)) is represented in a generalized form as:

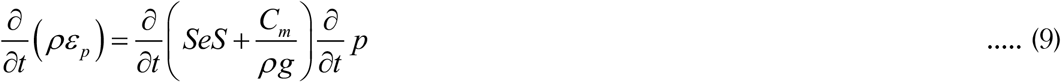

where, *ρε_p_* = ξ, *Se* = *S_n_*, *p* = φ, 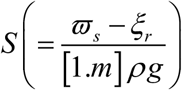 denotes the storage coefficient, which describes the variations in the storage of the fluid within the filter paper matrix, because of the expansion and compression of the void spaces and the liquid, when the matrix is fully saturated; 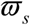 and *ξ_r_* represent the liquid volume fraction at saturation, and residual saturation respectively; 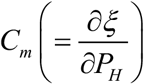 is the specific moisture which depicts the variation in the amount of moisture present in the sample, with the pressure head. (Buser, 2016; Rath et al., 2018; Rath and Toley, 2021) Within the Richard’s equation interface, the Darcy’s velocity is calculated as: 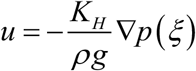, where, 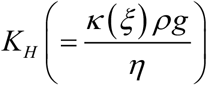 represents the hydraulic conductivity. Although the general form of the Richard’s equation in COMSOL does not contain an explicit variation in *ξ*, the use of van Genutchen parameters provides an implicit correlation between *φ* and *ξ*, thereby depicting the apposite physics. (Rath et al., 2018; Rath and Toley, 2021)

Subsequently, appropriate boundary and initial conditions were chosen, to solve the governing equation for fluid imbibition through the paper matrix. Since the entire set of experiments were performed under ambient conditions, it was natural to consider that the initial pressure within the porous media was close to the atmospheric pressure due to the slenderness of the paper devices. To put it into perspective, the length of the two devices were 20 and 2*5* mm (approx.), while the thickness of the filter paper was 0.18 mm. Hence the initial pressure was considered to be −101.325 [*kPa*], wherein the negative sign represents unsaturation. This value of the initial pressure was numerically close to the one considered by Chauhan et. al., (Chauhan and Toley, 2021) and Rath et. al., (Rath et al., 2018; Rath and Toley, 2021) who used the concept of an average capillary pressure. Additionally, to define the inlet boundary condition in our simulations for both the channels, we could not consider the capillary pressure value at saturation. (Hertaeg et al., 2019; Rath and Toley, 2021) This is due to the fact that, we used a *5* and 10 μL droplet of analyte at the deposition zone for the glucose and ketone sensors respectively, and we conjectured that this volume of water would be insufficient to fully saturate even the analyte deposition zone of either of the devices. Therefore, to validate this hypothesis, we calculated the value of normalized saturation 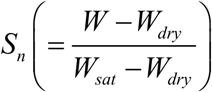, (Rath et al., 2018) where *W_dry_*, *W_sat_*, and *W* represent the weight of the device when fully dry, completely saturated, and partially saturated with either *5* or 10 μL droplet volume respectively. At least five individual glucose and ketone devices were carefully cut, and were heated at 100 °C for an hour, to eliminate any absorbed moisture. The devices were allowed to cool down to room temperature, and their weights were carefully measured using a sensitive balance, and this reading was marked as the dry weight. Subsequently, *5* and 10 μL droplets were placed on the glucose and AcAc device respectively, and their weights were measured after about 10 mins, and this value was denoted as *W*. Finally, all the devices were completely submerged in water for a period of 30 mins., and the surfaces were carefully patted with Kimwipes® and their weights measured, giving rise to *W_sat_*. We noticed appreciable variation in the value of normalized saturation not only between the two devices, but also between the values after depositing the specified liquid volumes, and those after the devices were fully saturated. Using this data, we used the Water Retention Curves (WRC), which relate the capillary pressure to water saturation in a porous matrix, as available in the literature for a Whatman grade-1 filter paper, (Buser, 2016; Chauhan and Toley, 2021; Rath et al., 2018; Rath and Toley, 2021) to arrive at the value of the inlet capillary pressure for the glucose (125 [*kPa*]) and ketone (140 [*kPa*]) sensors. However, it would be unrealistic to consider a constant value of inlet pressure throughout the length of the simulation time. Therefore, based on the variation of the capillary pressure with the normalized saturation (as per the WRC available in the literature (Buser, 2016; Rath et al., 2018)), the change in the capillary pressure with time was optimized to be consistent with the experimental results for both the devices, and that variation is expressed via the following expressions (equation 10 for the glucose device and 11 for the ketone sensor) :

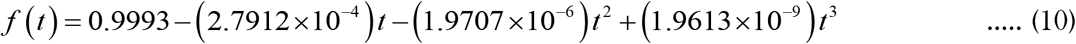

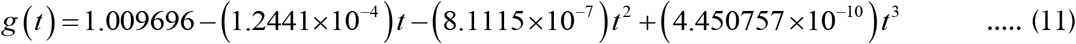

Finally, the capillary pressure value(s) obtained above (experimentally) were multiplied with equation (10) for glucose, and equation (11) for the ketone sensors, and used as non-homogeneous Dirichlet boundary condition(s) to describe the inflow of analyte into the two sensors respectively. Apart from the inlet condition, a non-flux Neumann boundary condition was used for all the other edges to represent the absence of fluid flow out of the devices (along the edges). The details of the various parameters used, the initial and boundary conditions for both the sensors is clearly depicted in Table S2 and Figures S2 and S3 of the ESI.

### 4.2. Species Transport in the Porous Matrix

One of the key aspects of modelling in our present work, is to study the transport and reaction amongst various species within the porous media. To accomplish this objective, we coupled the Richards’ equation with the species transport equation, which for a porous substrate is given by (Rath and Toley, 2021):

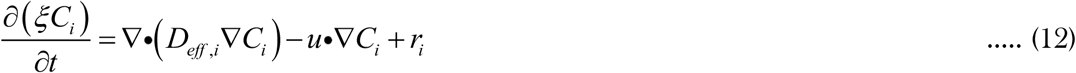

where, *C_i_* denotes the concentration of species *i* in the system, *D_eff, i_* denotes the effective diffusivity of the *i^th^* species in the porous matrix, represented by the Millington and Quirk model (Millington, 1959; Millington and Quirk, 1959), *u* is the Darcy’s velocity, and *r_i_* represents the rate of reaction pertaining to the species *i*, the equations of which would be described in the subsequent sub-section for the two assays. It is to be noted that the species transport equation for a porous media consists of the term *ξ*, which as specified previously, denotes the volume fraction of the fluid within the porous matrix, and such a term is absent for the case of a non-porous material. (Leal, 2007; William, 1998)

An interesting point to be noted here is the coupling between the two modules. Specifically, this is a classic one-way coupling of the governing equations, wherein the solution to the Richards’ equation is independent of the species transport, whereas the latter strongly depends on the flow-field calculated using the Richards’ equation interface. To effectuate this coupling within the COMSOL software, we used the “partially saturated porous media” feature of the “Transport of dilute species” module. The initial concentrations of all the species pertaining to the two assays were used in the respective domains (reagent zone and deposition zone), in accordance with our experiments (see section S3 of the ESI). Additionally, the inlet boundary condition was set as a logical constraint based on the solution of the Richard’s equation. The rationale used was that, the analyte would be carried into the porous matrix along with the fluid, and in the expanse un-imbibed by the fluid, the concentration of the analyte would be zero. (Buser, 2016; Rath and Toley, 2021) Therefore, the following logical condition was used to denote the inlet concentrations for the two assays:

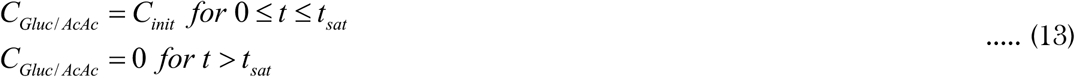

where, *C_Gluc/ AcAc_* denotes the concentration of glucose or AcAc, *C_init_* refers to the individual value pertaining to the concentration range used in the experiments for the two assays, i.e., *C_init_* ranges from 1 to 1*5* 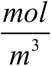 and 1 to 10 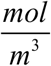 for the glucose and ketone assays respectively; *t_sat_* refers to the time taken by the fluid to fully saturate the porous filter paper. Based on the standalone numerical results of the Richard’s equation, the value of *t_sat_* was 320 s for the glucose device, and 850 s for the ketone device. Subsequently, within the “partially saturated porous media” interface, the values of the porosity, the specific moisture capacity, and liquid volume fraction were selected based on the solution of the Richard’s equation. Additionally, the time-varying pressure head was chosen to represent the temporal change in the fluid fraction. Finally, the abovementioned logical construct was used as a Dirichlet boundary condition to describe the species inlet at the deposition zone, while a homogeneous (no-flux) Neumann boundary condition was used on all the edges of the two devices to denote the absence of outflow for any of the species (see section S3 of the ESI). The coupled equations were solved for a total duration of 700 s and 12*5*0 s, with a time-step of 10 s, to simulate the glucose and ketone devices respectively.

### 4.3. Kinetics of the Glucose Assay

The enzymatic oxidation of glucose has long been used as a reliable mechanism to determine the glucose content in biofluids. (Cao et al., 2020; Chaiyo et al., 2018; Chauhan and Toley, 2021; Chen et al., 2012, 2020; Gu et al., 2019; Kap et al., 2021; Khan et al., 2019; Kim et al., 2020; Klasner et al., 2010; Leitner et al., 2001; Liu et al., 2019; Ngo et al., 2021; Rossini; et al., 2021; Rossini et al., 2019; Singh et al., 2021; Tam et al., 2021; Zhu et al., 2014) The compatibility of this reaction with paper devices has been one of the major reasons for the wide-spread applicability and potential commercialization of μPADs in healthcare monitoring of various ailments in general, and diabetes in particular. (Klasner et al., 2010; Martinez et al., 2010b, 2010a; Nishat et al., 2021; Selvakumar and Kathiravan, 2021; Tseng et al., 2021; Yetisen et al., 2013) One of the unique aspects of this assay is the coupled kinetics involving two enzyme-catalyzed reactions. (Bateman and Evans, 1995) The first reaction occurs between glucose (analyte), and the enzyme glucose oxidase, which is a Flavin adenine dinucleotide (FAD)-dependent oxidoreductase, and has a very high affinity to oxidize glucose in comparison with other sugars. (Bateman and Evans, 1995; Cao et al., 2020; Leitner et al., 2001; Nery; and Kubota, 2016; Rossini et al., 2019; Tao et al., 2009) This reaction in presence of oxygen, gives rise to hydrogen peroxide, and the scheme is represented as follows (Tao et al., 2009):

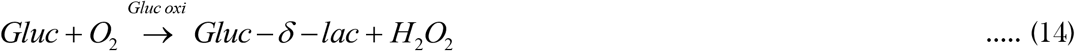

where, *Gluc* represents glucose, *Gluc oxi* is glucose oxidase, and *Gluc*–*δ*–*lac* denotes glucono–*δ*–*lactone*.

In an aqueous solution, the produced glucono-*δ* -lactone, reacts further with water, giving rise to gluconic acid. (Bateman and Evans, 1995; Tao et al., 2009) Additionally, it is worth noting that the enzyme glucose oxidase is uniquely specific towards β-D-glucose, and does not catalyse its muta-rotated form α-D-glucose. (Tao et al., 2009) However, in a solution these two forms of glucose co-exist, and the interconversion between these two isomers is a very slow reaction. Therefore, we assume the existence of an equilibrium mixture between these two forms of glucose, and given that only the β-D-glucose form reacts with the enzyme, the above reaction scheme represented by equation (14) would be valid. (Bateman and Evans, 1995; Leitner et al., 2001; Tao et al., 2009) Broadly, the enzyme oxidizes glucose to glucono-*δ* -lactone, and the oxygen reacts with the reduced form of the enzyme to re-oxidize it, thereby finally producing hydrogen peroxide in the process. (Bateman and Evans, 1995; Ngo et al., 2021; Tao et al., 2009) The entire kinetics could be approximated satisfactorily using the Michaelis-Menten kinetics, and the rate of formation of *H*_2_*O*_2_ is given by the following equation (Bateman and Evans, 1995; Leitner et al., 2001; Singh et al., 2021):

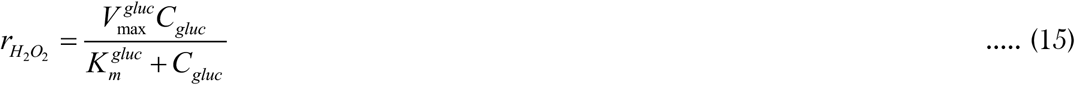

where, 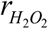 denotes the initial rate of formation of the product pertaining to the oxidation of glucose, 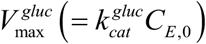 is the maximum value of the initial reaction rate at high substrate concentration, 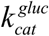 is the rate constant associated with the enzyme, *C_E,0_* is the concentration of the enzyme, 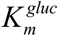 is the concentration of the substrate (glucose) when 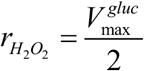, and it is an indicator of the enzyme performance (higher values of 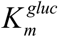 indicate lower enzymatic performance). The values of 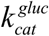, and 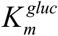 for the glucose oxidation reaction are available in the literature. (Chen et al., 2012; Leitner et al., 2001; Singh et al., 2021; Tao et al., 2009)

The hydrogen peroxide produced in the abovementioned reaction, would now react with potassium iodide in another enzyme-catalyzed reaction, in presence of horseradish peroxidase. (Bjorksten, 1970; Dolman et al., 1975; Dunford and Stillman, 1976; Ozaki and de Montellano, 1995; Roman; and Dunford, 1972; Ugarova; et al., 1981) The reaction is conjectured to proceed via the scheme depicted below (Dolman et al., 1975; Dunford and Stillman, 1976; Ugarova; et al., 1981):

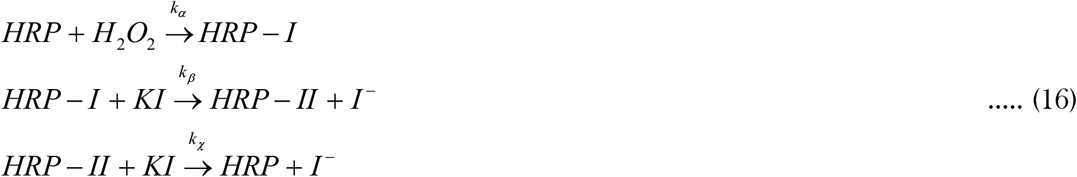

where, *HRP* denotes horseradish peroxidase, *HRP* – *I* and *HRP* – *II* are the reduced form of the enzymes (intermediate products), and *KI* denotes potassium iodide.

The initial rate of formation of *I*^−^ for the two-substrate, enzyme catalyzed reaction is given as follows (Dunford and Stillman, 1976; Ugarova; et al., 1981):

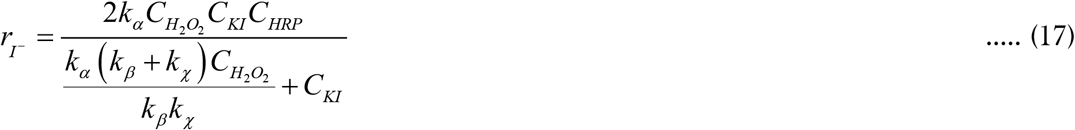

We consider that the above kinetics could be sufficiently represented by the Michaelis-Menten equation, resulting in the following variables (Ugarova; et al., 1981):

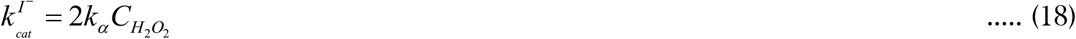

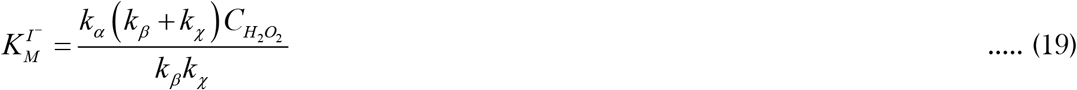

The values of 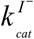 and 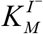 for the HRP catalyzed oxidation of hydrogen peroxide is available in the literature. (Bjorksten, 1970; Dolman et al., 1975; Dunford and Stillman, 1976; Ozaki and de Montellano, 1995; Roman; and Dunford, 1972; Ugarova; et al., 1981)

To demonstrate the glucose assay kinetics in COMSOL Multiphysics (v. *5*.4), the concentration of glucose was varied from 1 to 1*5* 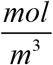, while those of glucose oxidase, HRP, and KI were 0.01*5*, 0.00376, and 600 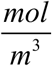 respectively (taken in accordance with the experiments). The inlet boundary condition for glucose was placed at the analyte deposition zone, while those for HRP, KI, and glucose oxidase were placed at the reagent deposition zone. The reaction kinetics were considered throughout the reagent deposition zone and the detection zone, as depicted in Figure S2 of the ESI. A free triangular mesh was chosen for the simulation, consisting of 1069 elements (including domain and boundary), and a total of 11957 degrees of freedom were solved to obtain the results.

### 4.4. Kinetics of the AcAc Assay

One of the key differences between the kinetics describing the glucose and ketone assays, lies in the nature of the reaction itself. The transformation of D-glucose to produce the final-colored product involves two simultaneous enzyme-catalysed reactions, whereas, the AcAc assay mechanism, is non-catalytic and can be described using pseudo-first order kinetics. (Laios et al., 1986; Laios and Pardue, 1993) The nitroprusside test, as it is commonly known, consists of a reaction amongst acetoacetate, glycine, and sodium nitroprusside giving rise to a magenta-coloured product (absorption maxima at *5*40 nm). (Laios and Pardue, 1993) This system of reactions is conjectured to progress in three steps.(Laios et al., 1986; Laios and Pardue, 1993) Firstly, the acetoacetate reacts with glycine to produce enamine. Secondly, the product of the first step reacts with sodium nitroprusside to from a coloured product which gradually decays into a stable product, in the third step. (Laios and Pardue, 1993) The reaction scheme is depicted in equation (20). (Laios and Pardue, 1993)

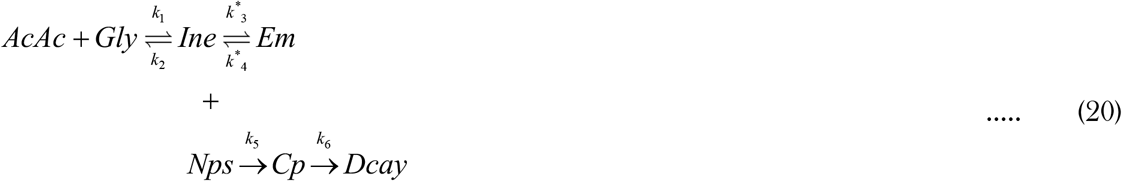

Where, *AcAc* denotes acetoacetate, *Gly* represents glycine, *Ine* is the imine derivative, *Em* depicts the enamine, *Nps* is sodium nitroprusside, *Cp* denotes the colored product, and *Dcay* is the product of the decay reaction. The reaction rate constants represented by *k*_1_, *k*_2_, *k*^*^_3_, *k*^*^_4_, *k*_5_, and *k*_6_, are available in the literature. (Laios and Pardue, 1993)

Although the proposed reaction mechanism could be described using equation (20), experimental examinations reported in the literature, clearly indicate that the reaction between glycine and acetoacetate follows a pseudo-first order kinetics. (Laios et al., 1986; Laios and Pardue, 1993; Molina et al., 1989) Therefore, in accordance with our experiments, the concentration of glycine is several times higher than that of AcAc, and hence there is no perceptible change in the glycine concentration as the reaction proceeds. Additionally, the tautomerization of imine to enamine involving the rate constants *k*^*^_3_ and *k*^*^_4_ could be safely neglected owing to the fact that the formation of the imine derivative is the rate limiting step. (Laios and Pardue, 1993) Subsequently, the imine produced in the first step reacts with sodium nitroprusside rapidly, following another pseudo-first order kinetics, resulting in the formation of the coloured product which has a tendency to decay, and form a stable product. (Laios et al., 1986; Laios and Pardue, 1993) However, herein the decay reaction (step 3 in the abovementioned scheme) was not considered due to the fact that the product of step two which has an absorption maximum at *5*40 nm, was the one that was detected experimentally, owing to its colour band in the visible spectrum. (Klasner et al., 2010; Laios and Pardue, 1993) With these considerations, the rate equation for acetoacetate, the imine derivative, and the coloured product are given by the following equations (Laios et al., 1986; Laios and Pardue, 1993):

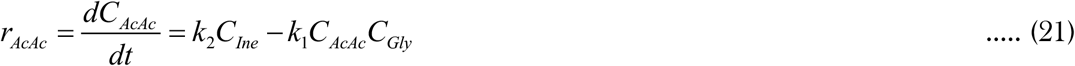

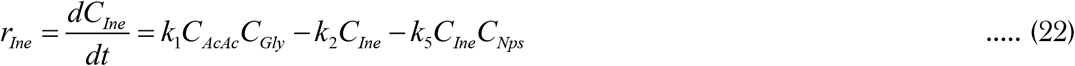

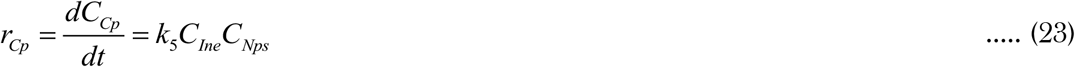

Within the framework of the numerical simulation, we varied the initial concentration of AcAc from 1 to 10 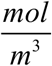, while the concentration of glycine and sodium nitroprusside were 100 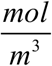, and 176 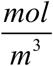, respectively (in accordance with the experiments). The boundary conditions pertaining to the inlet concentration of AcAc was placed in the analyte zone, and those for glycine and nitroprusside were in the first and second reagent zone respectively, as depicted in Figure S3 of the ESI. Additionally, it could be possible that the flow of the analyte carries with it, some of the reagents, and hence the reaction domains were chosen such that, both the reagent zones, the narrow flow-path connecting them, and the detection zone, could all serve as reaction media for the first reaction (equations 21 and 22). Whereas, the detection zone, and the second reagent zone would only be used to effectuate the kinetic of the second reaction (equations 22 and 23). Although the pre-defined meshes could be used for the simulation, we chose a free triangular mesh for its versatility. The final mesh size was optimised such that it consisted of 1429 elements (domain and boundary), and a total of 24733 degrees of freedom were solved to obtain the results of the numerical simulation pertaining to the AcAc assay.

## 5. Results and Discussion

In the present study, the emphasis was initially placed on the successful fabrication of the two sensors capable of accurately estimating the amount of glucose and ketone in a given urine sample. However, as would be elucidated subsequently, the experimental visualization of the rate and zone of coloration were intriguing owing to the fact that the colored product gradually accumulated in a small, curved expanse at the edge of the detection zone. Therefore, it was imperative to comprehend the physics of the process that would aid in designing better and efficient sensors for rapid and accurate analysis. But there was a significant paucity in literature pertaining to the theoretical or numerical investigation of flow coupled reaction in a paper matrix. Hence, as an earnest attempt to bridge that gap, and to understand the pertinent physics, a numerical investigation of the flow field and the reaction kinetics was undertaken. The following sub-sections present the details of the numerical simulation along with the experimentally observed variation in the concentration of the colored product for both the sensors.

### 5.1. Flow Through the Paper Matrix

The results of the time-varying fluid imbibition through the porous filter paper obtained by solving the Richards’ equation in COMSOL, for both the devices is depicted in Figures 4 and 5. The snapshots presented in Figures 4 (a) and 5 (a) correspond to the temporal variation in the liquid volume fraction, while those portrayed in Figures 4 (b) and 5 (b) represent the velocity streamlines (Darcy’s velocity) and pressure contours corresponding to the fluid imbibition. It is clear from the figures that the liquid saturated the glucose device within 320 s, while it takes about 8*5*0 s for the analyte to traverse from one end to the other in the ketone device. The difference between the time taken is not only due to the variation in the lengths of the two devices, but also owing to the presence of two converging-diverging sections in the AcAc device, that demands greater pressure drop for the same amount of fluid flow. The detailed spatio-temporal variation in the liquid volume fraction along the flow path for the two devices is presented in the supplemental video SV1. As observed from SV1, and also from the snapshots presented in Figure *5*(a), the fluid imbibes the first reagent zone of the ketone sensor within 210 s, while it takes an additional 400 s (approx.) to saturate the second zone. Another reason for the time-lag is the time-varying inlet pressure variation, which is in-line with the experiments.

**Figure 4:**
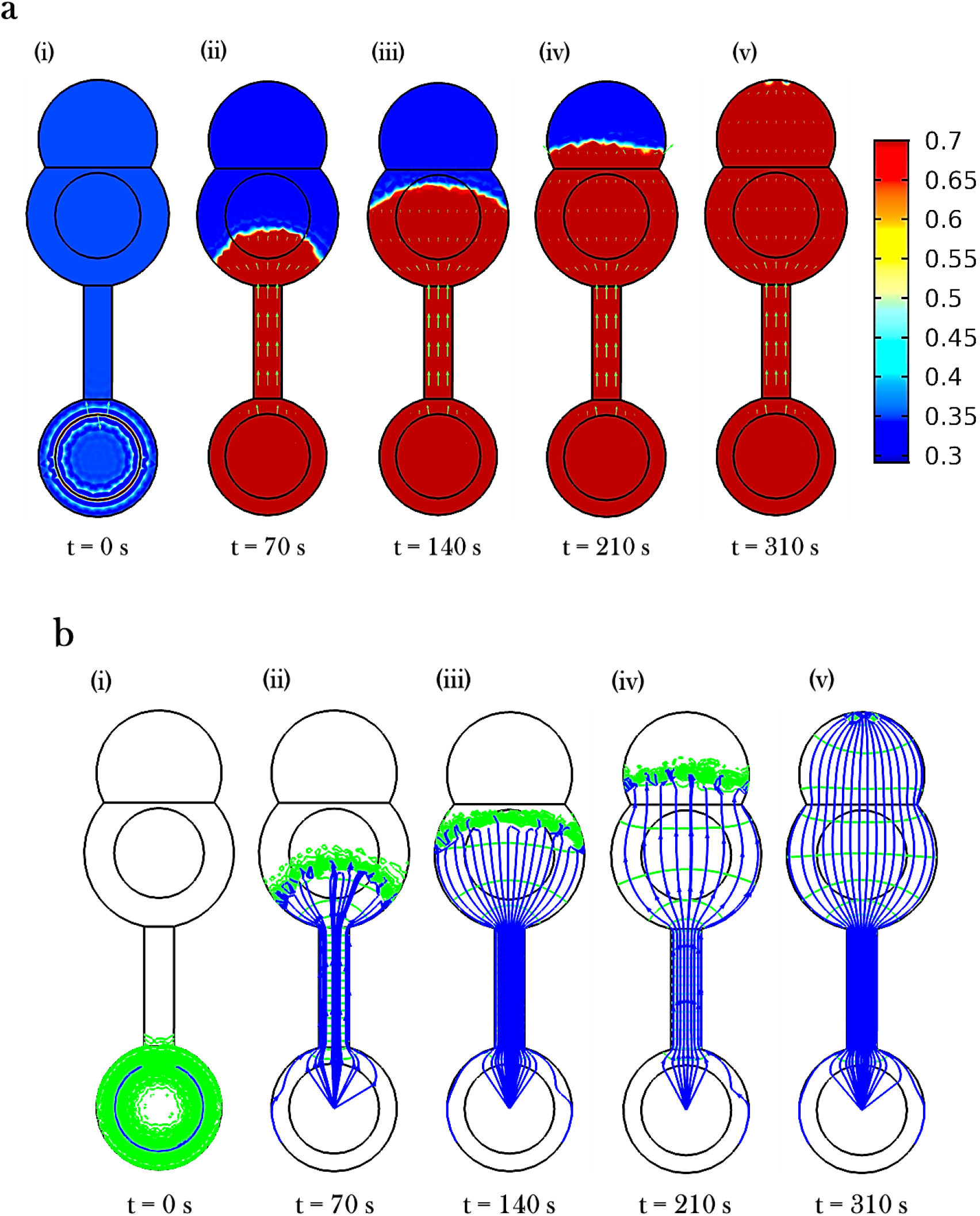
(a) Numerical simulation results of the spatio-temporal variation in the liquid volume fraction along the flow-path for the glucose sensor. (b) Pressure contours and Darcy’s velocity streamlines for the time-varying fluid imbibition in the glucose device.

**Figure 5:**
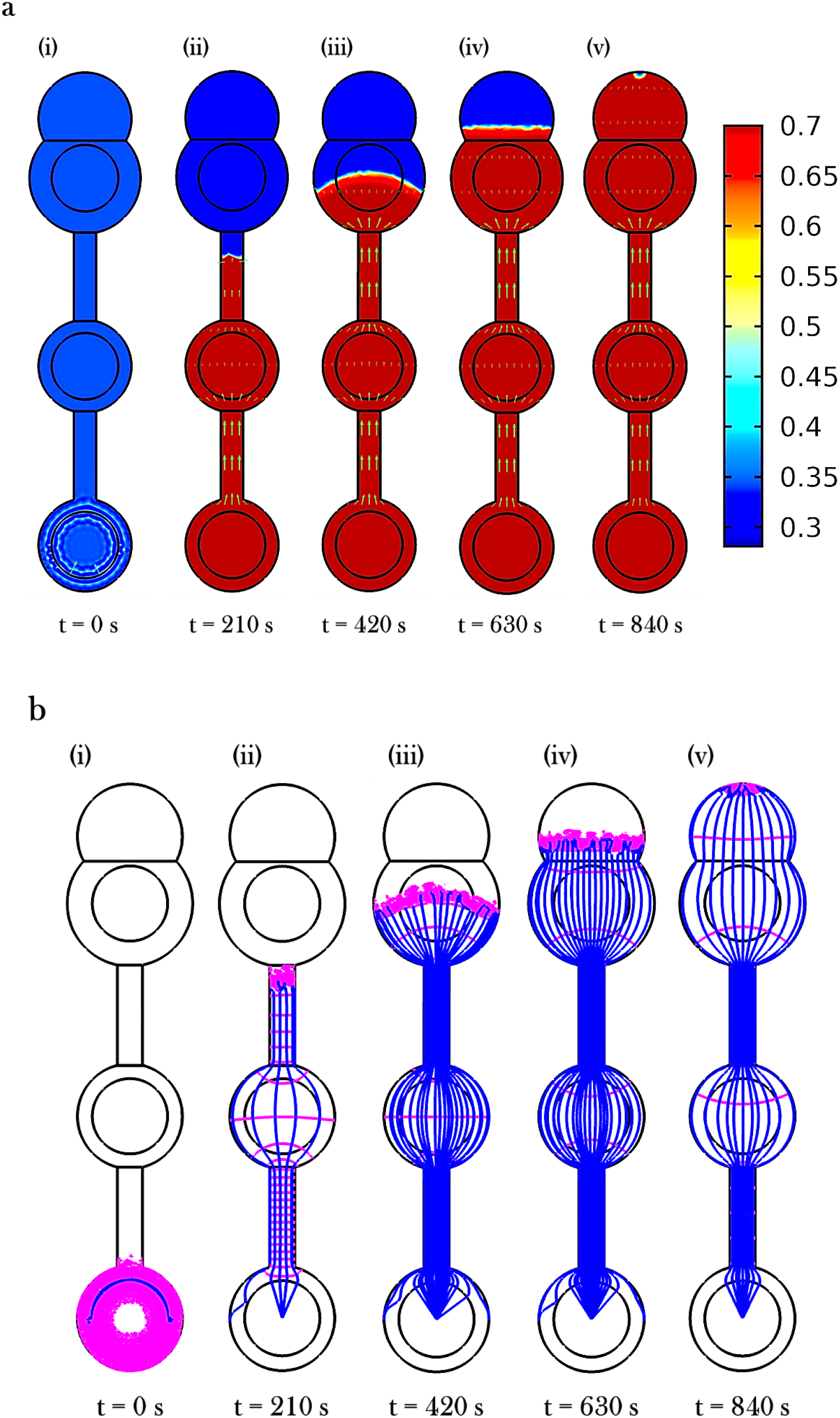
(a) Results of the numerical simulation pertaining to the spatio-temporal variation in the liquid volume fraction along the flow-path for the ketone sensor. (b) Pressure contours and Darcy’s velocity streamlines for the time-varying liquid imbibition in the ketone device.

We note here that, a detailed experimental investigation of the water retention curve pertaining to the device geometry, could shed additional light on the variation of the capillary pressure with saturation. However, such a study comparing the numerical and experimental flow characteristics coupled with understanding the kinetics of the reactions on a paper device, is a work in progress, and is a part of the future work. We wish to highlight that the present study outlines the detailed methodology to effectively comprehend the flow-induced kinetics in a paper device with variable cross-section, while considering a specific volume of inlet liquid through the devices, as opposed to a multitude of studies that consider infinite volume of liquid at the source.(Modha et al., 2021; Rath et al., 2018) Another unique feature of our work, lies in demonstrating the simplicity with which the flow through a variable cross-section paper device could be modelled, using the Richards’ equation in COMSOL Multiphysics®, which offers a favourable solution to the limitations of Lucas-Washburn equation for modelling a 2D flow in a paper-device.(Buser, 2016; Rath et al., 2018)

### 5.2. Glucose Assay

The glucose assay, as stated previously, was performed for the concentration range of 1 mM to 1*5* mM and the resultant intensity values (dimensionless) as a function of the glucose content in the urine sample are represented in Figure 6 (a). It is to be underscored that the acquired intensity values are divided by the maximum intensity (white region of the filter paper) for a particular image to rule out the errors that could have emanated from the differences in the initial settings of the microscope.

**Figure 6:**
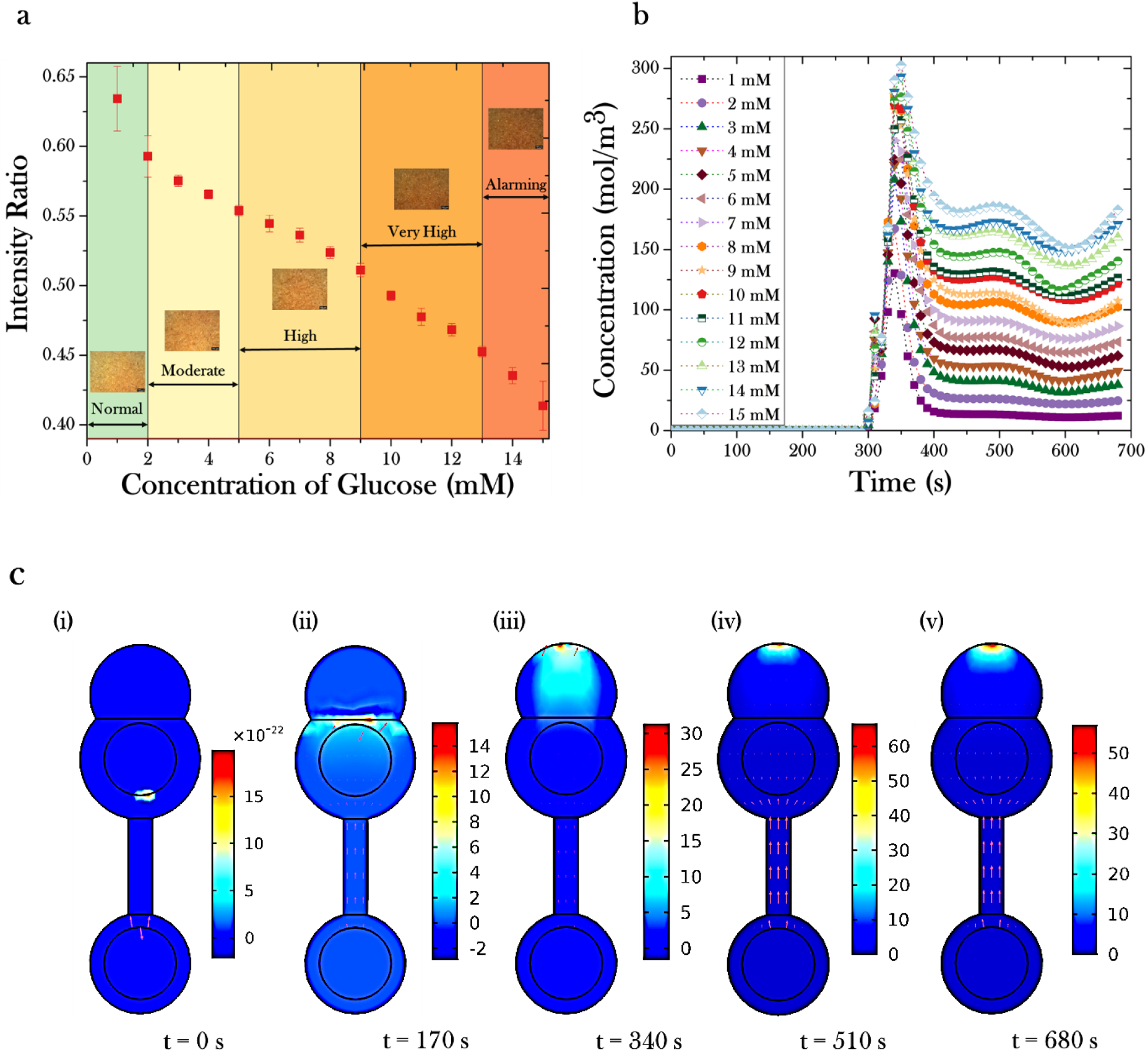
(a) Phase diagram pertaining to the experimental variation in the intensity ratio of the colored product with change in the glucose concentration. The insets are representative images of the detection zone in the glucose sensor corresponding to the concentration bands. A clear decrease in the gray values could be observed in the images, with an increase in the concentration of the glucose in urine. The phase-diagram distinctly delineates the different bands relative to the health-status (amount of glucose in urine) of a subject. (b) Simulation results of the temporal variation in the instantaneous concentration of the colored product in the glucose sensor, measured at the edge of the detection zone. The flow-induced enzyme-catalysed reaction triggers an initial rise followed by gradual equilibration of the colored product concentration. (c) Snapshots of the spatio-temporal variation in the formation of the colored product in the glucose sensor, obtained from the numerical simulation. The results correspond to a glucose concentration of *5* mM.

To determine the intensity ratio ranges for classifying the samples from normal to alarming contents of the glucose, 99% confidence interval for all the concentrations of all tested samples was calculated using the following equation:

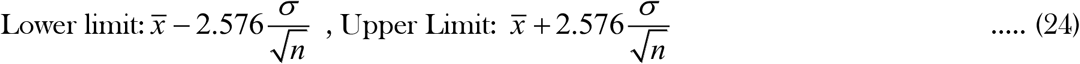

where, 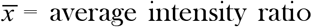, σ = standard deviation and n = number of samples. The parameters used for estimating the 99% confidence interval are shown in Table S3 of ESI. The intensity ratio ranges have been correlated with the indicator ranges of glucose and have been depicted in Table 1 and Figure 6 (a). The intensity ratio range, presented herewith can serve as a potential biomarker to determine the approximate glucose content of an unknown urine.

**Table 1:** Intensity ratio ranges for detecting the glucose content

| S. No. | Concentration of glucose, C<br>(mM) | Intensity ratio (I) | Diagnosis |
| --- | --- | --- | --- |
| 1 | $C \leq 2$ | $I \geq 0.58$ | Normal |
| 2 | $2 < C \leq 5$ | $0.55 \leq I < 0.58$ | Moderate |
| 3 | $5 < C \leq 9$ | $0.51 \leq I < 0.55$ | High |
| 4 | $9 < C \leq 13$ | $0.45 \leq I < 0.51$ | Very High |
| 5 | $C > 13$ | $I < 0.45$ | Alarming |

Additionally, the flow induced reaction between glucose and the different reagents results in the formation of a golden-brown colored product, as stated previously. The variation in the rate of formation of the colored product for three different glucose concentrations (viz. 1, 8, and 1*5* mM) is depicted in the supplemental video SV2, while the instantaneous concentration of the colored product for all the different initial glucose concentrations considered in our experimental investigation is depicted in Figure 6(b). The snapshots presented in Figure 6(c) correspond to the rate of final product formation for a glucose concentration of *5* mM. As observed from the figure, the color product begins to form, as soon as the analyte comes in contact with the reagent. The rate of formation of hydrogen peroxide and the color product are in accordance with the kinetics described in the previous section (see section 4.3). The design of the glucose device provides an optimum flow-path for the analyte to come into contact with the respective reagents. The formed color product is gradually carried along with the flow, and accumulates in the expanse dedicated for imaging the final product as observed from the experiments, thereby validating the results of the numerical simulations.

### 5.3. Ketone assay

The AcAc assay was performed for the concentration range of 1 mM to 10 mM and the resultant intensity ratios (dimensionless) are plotted against the ketone content in urine, as shown in Figure 7 (a).

**Figure 7:**
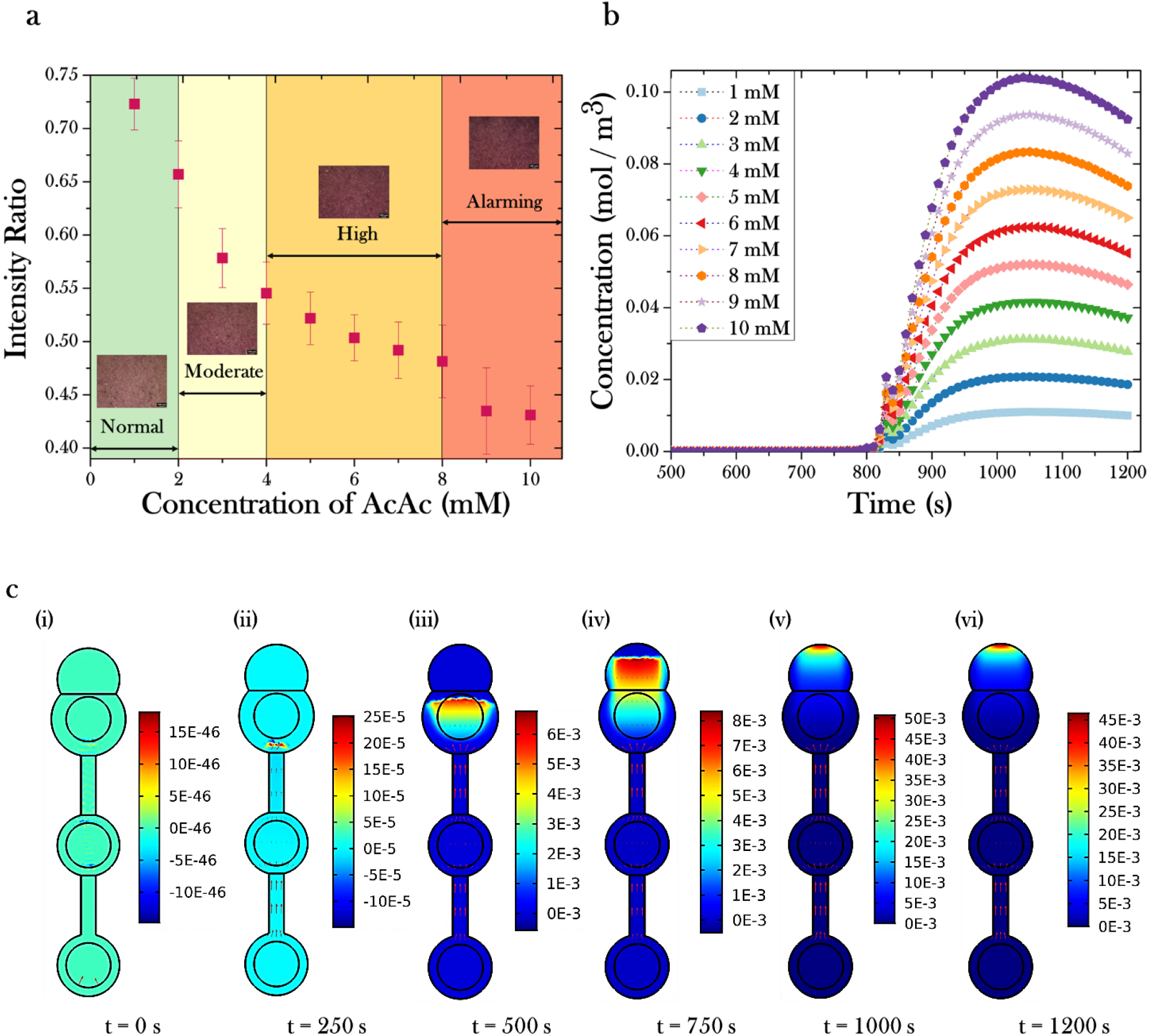
(a) Phase diagram corresponding to the experimental variation in the intensity ratio of the colored product with change in the AcAc content in the urine sample. The insets are representative images of the detection zone in the ketone sensor in relation to the concentration bands. A distinct decrease in the image intensity as a function of increasing AcAc content could be observed in the images which correspond to the increased amount of colored product formation, thereby causing a darker color in the detection zone. The phase-diagram clearly delineates the different bands relative to the health-status (amount of AcAc in urine) of a subject. (b) Simulation results of the temporal variation in the instantaneous concentration of the colored product in the ketone sensor, measured at the edge of the detection zone. The flow-induced non-catalysed reaction occurs in two separate zones, following which the colored product is gradually accumulated at the edge of the detection zone. (c) Snapshots of the spatio-temporal variation in the formation of the colored product in the ketone sensor, obtained from the numerical simulation. The results correspond to a AcAc concentration of *5* mM.

As elucidated in the previous sub-section, a 99% confidence interval for all the concentrations (shown in Table S4 of ESI) of samples were calculated to correlate the intensity ratios with the indicator ranges of ketone (as shown in Table 2 and Figure 7 (a)).

**Table 2:**
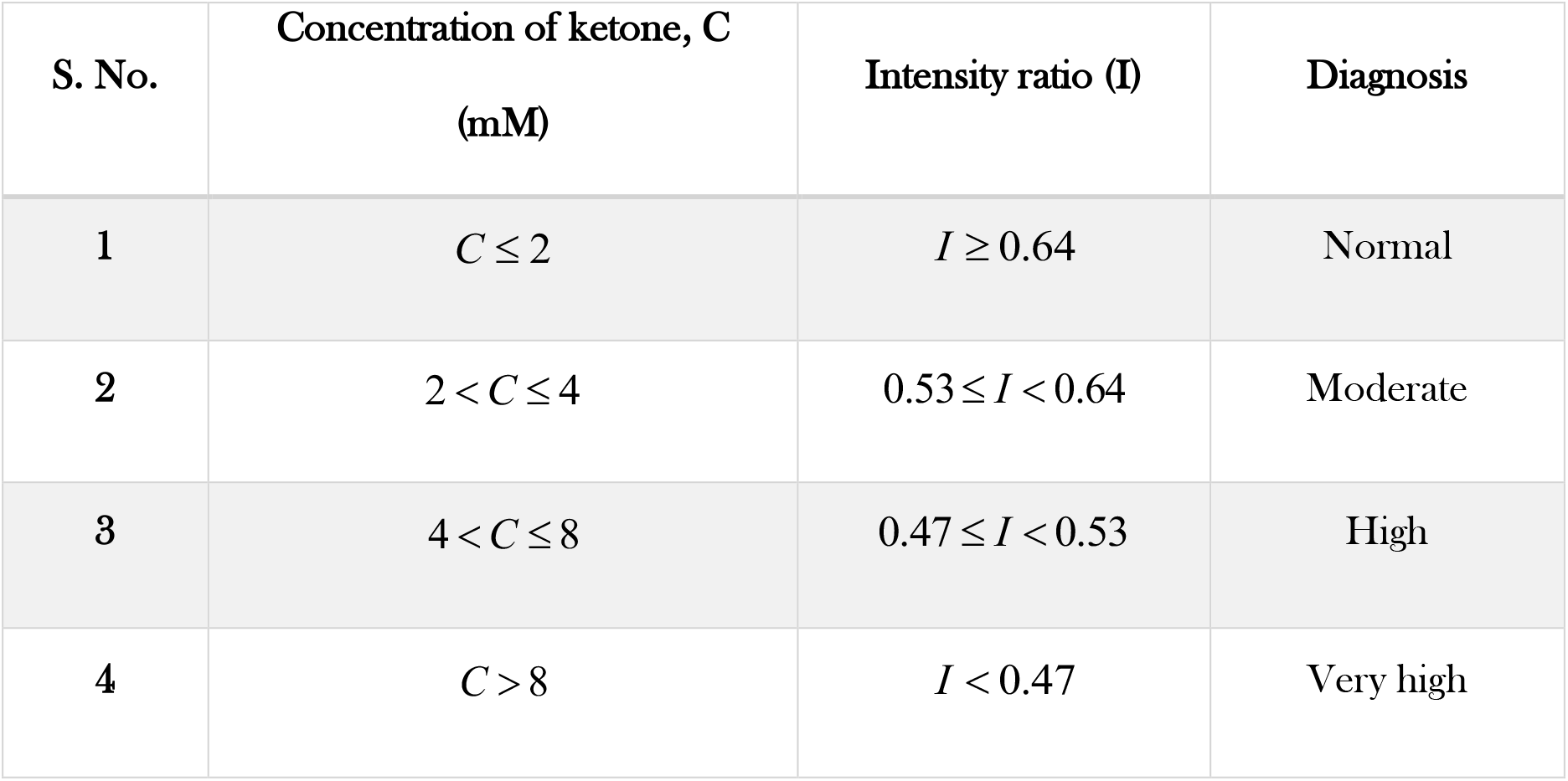
Intensity ratio ranges for detecting the ketone content

While the intensity ratios are depicted in Figure 7(a), the time varying concentration of the final-colored product for different initial concentrations of acetoacetate in urine is presented in Figure 7(b). The effect of flow on the rate of product formation for three initial AcAc concentrations (viz. 1, *5*, and 10 mM) is elucidated in the supplemental video SV3, and the snapshot presented in Figure 7(c) are for *5* mM AcAc concentration. As observed from the figure (Figure 7(c)), the imine derivative produced in the first reagent deposition zone, is gradually carried along with the analyte (urine) into the second zone, wherein it reacts with nitroprusside, giving rise to a deep-violet colored compound, which has an absorption maximum in the visible spectrum. The concentration of the final product increases with the initial concentration of the ketone present in the analyte, as observed from the intensity curve obtained experimentally, thereby corroborating the numerical results. It is well overt that the ketone moieties produced by the body, in the liver are used as sources of energy, when the glucose content is gradually depleted. (Khan et al., 2004; Klasner et al., 2010; Laffel, 1999) Therefore, the two main ketones generated in this process are 3-β-hydroxybutyrate (BHB) and acetoacetate, while a minor quantity of acetone is also produced in the process, but it is least abundant, making the detection of acetone difficult and unreliable for diabetes detection.(Khan et al., 2004; Klasner et al., 2010; Klocker et al., 2013; Laffel, 1999) Although a combined estimation of BHB and AcAc would provide a complete picture of the ketone content in a subject, we noticed that there were significant stability issues pertaining to the enzyme used in the BHB assay, (Wang et al., 2016) especially over a paper substrate. Therefore, we intend to develop an alternative methodology for the characterization of BHB in biofluids on a paper device, as a part of the future work, along with the experimental determination of the flow-field, and imbibition. Additionally, we would also like to point out that, we have exercised utmost caution in perusing and choosing the relevant kinetic equations for the two assays (in-line with our experimental conditions), as per the available literature. However, due to the complex nature of the system, the results presented here could be used as a qualitative estimate of the concentration profile, while a detailed quantitative study is out of scope of the present work, and would be undertaken in the future. We would still like to emphasize on the fact that the present numerical work, is a first of its kind study of the flow modulated chemical kinetics within a paper matrix, and would serve as an initial estimate for obtaining a clear view on the various design parameters essential for successfully fabricating novel paper-based microfluidic devices.

## 6. In-house App. for Rapid Glucose and Ketone Detection

Although extreme caution was exercised while analyzing the images using the open-source software – “ImageJ (v.2.0 Fiji)”, there is always a possibility albeit miniscule, of introducing inadvertent human errors into the analysis. Therefore, it is imperative to perform the image investigation with minimal human intervention. To that end, an automated image analysis subroutine was developed in-house, and later transformed to a desktop and web application, viz. Rapid Glucose and Ketone Detector (RGAD), using MATLAB (v. 2020b). The graphical user interface (GUI) of the desktop app. is depicted in Figure 8. Within the app., the user is initially required to upload a standard white image as a reference to the ambient lighting conditions, since the perception of image intensity is an intrinsic function of the background brightness. Subsequently, the glucose and ketone specific images are also to be uploaded. Post transferring the three images into the software, the image analysis algorithm, is activated, wherein the three images are cropped to a predefined rectangular expanse to ensure the elimination of any uneven background. The images are then processed such that the mean intensity of all the pixels within the cropped region are calculated for all the three pictures. Thereafter, the average intensity of the glucose and ketone images are divided by the mean intensity of the standard image, resulting in the respective intensity ratio. The values thus obtained are compared with the standard calibration curve (obtained experimentally, as described in the previous section) to estimate the relative color grade and category (normal or otherwise) based on the glucose and ketone content of the samples. The representative usage of the RGAD app. at different background light-settings is depicted in supplemental video, viz. SV 4. It is to be noted that the results obtained from the MATLAB application were extremely repeatable and reliable, since the difference in the intensity ratios between ImageJ and the RGAD app. was negligible (0.01-0.4*5*%) (as shown in Table S5 and S6 of the ESI), thereby underscoring the effectiveness of the developed application.

**Figure 8:**
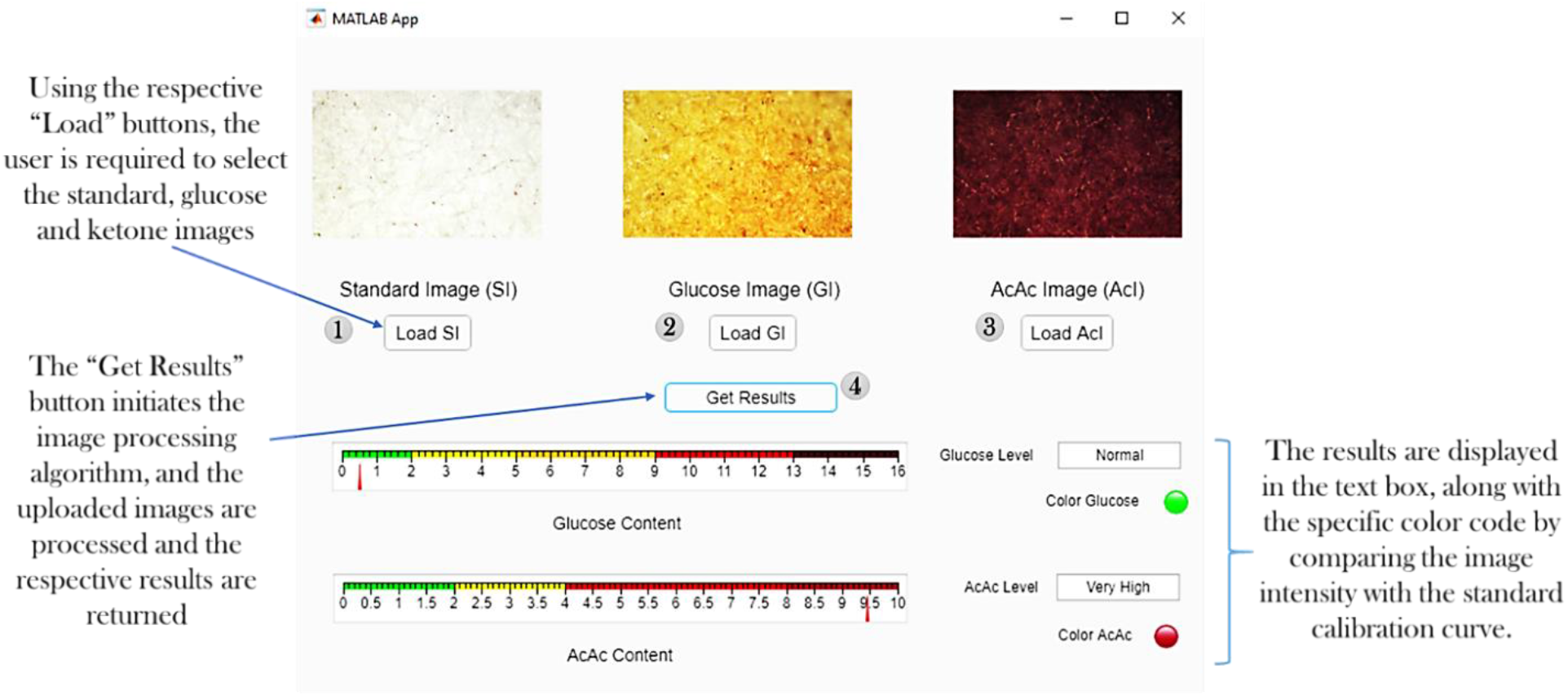
Schematic depicting the graphical user interface of the in-house developed MATLAB application (RGAD). The user is initially required to upload three images corresponding to the standard white, glucose, and ketone. The image analyzing pipelines processes the three images, computes the intensity ratio, and displays the corresponding result category (normal or otherwise) in the text box, along with the relevant color code. A pointer and a slider with the color scheme corresponding to the standardization curve are also depicted in the app. Depending on the calculated intensity ratio, the pointers slide, and display a numeric value corresponding to the approximate glucose and/or ketone content in the sample.

## 7. Conclusion

Herein, we present a facile laser-printing based methodology for fabricating μPADs capable of simultaneously detecting glucose and acetoacetate in urine. The detailed strategy pertaining to the choice of the filter paper and the optimization of the device outline is also presented, along with the rationale for selecting the final design for the two sensors. The glucose device consisted of an analyte zone and a single reagent deposition zone, while the AcAc device was fabricated with two reagent areas in accordance with the kinetics, to facilitate the accumulation of the final-coloured product at a specific expanse (detection zone) to aid in further analysis. The analyte (urine), upon being deposited, wicked through the paper-device and gave rise to a golden-brown coloured product (for glucose), and a deep-violet coloured product (for ketone), which was further captured using a camera connected to an upright microscope. The obtained images were analysed using an open-source software, resulting in an intensity vs. concentration plot, which was used for the creation of a standardization curve pertaining to the two assays. Additionally, the flow mediated enzyme-catalysed reaction (for glucose), and the non-catalysed reactions (for AcAc) in a paper device was modelled and simulated using COMSOL Multiphysics. A good agreement was observed between the experiments and numerical simulations corresponding to the coloration zones of the final product in both the sensors, thereby corroborating the model. Finally, to widen the scope of our present work, we also developed a web and desktop-based application, which was tested at different light settings. We observed a negligible variation (less than 0.*5*%) between the results descried from the in-house developed app., and the open-source software (ImageJ, Fiji v 2.0). We believe that the present numerical work, and the facile laser-printing based method could serve as reliable tools for the design optimization and low-cost fabrication of paper-based devices, thereby augmenting their widespread applicability and facilitating analyte detection in resource limited settings.

## Supporting information

Supplementary Section for the Manuscript

## Author Contributions

**Manikuntala Mukhopadhyay**: Conceptualization, Data curation, Formal analysis, Investigation, Visualization, Methodology, Writing – Original draft; **Sri Ganesh Subramanian**: Data curation, Formal analysis, Investigation, Methodology, Software, Visualization, Writing – Original draft; **K. Vijaya Durga**: Data curation, Methodology, Investigation; **Debashish Sarkar**: Investigation, Supervision, Writing – review & editing; **Sunando DasGupta**: Investigation, Project administration, Resources, Funding acquisition, Supervision, Writing – review & editing

## Acknowledgement

One of the authors, Manikuntala Mukhopadhyay, gratefully acknowledges the funding and support received from BIRAC-SRISTI, through the Gandhian Young Technological Innovation Award, 2019.The authors also gratefully acknowledge the financial support provided by the Indian Institute of Technology Kharagpur, India [Sanction Letter no.: <u>IIT/SRIC/ATDC/CEM/2013-14/118</u>, dated 19.12.2013].

## References

Abe, K., Suzuki, K., Citterio, D., 2008. Inkjet-printed microfluidic multianalyte chemical sensing paper. Anal. Chem. 80, 6928–6934. https://doi.org/10.1021/ac800604v

Ahmed, S., Bui, M.-P.N., Abbas, A., 2016. Paper-based chemical and biological sensors: Engineering aspects. Biosens. Bioelectron. 77, 249–263.

Aksorn, J., Teepoo, S., 2020. Development of the simultaneous colorimetric enzymatic detection of sucrose, fructose and glucose using a microfluidic paper-based analytical device. Talanta 207, 120302 (1-8). https://doi.org/10.1016/j.talanta.2019.120302

Akyazi, T., Basabe-Desmonts, L., Benito-Lopez, F., 2018. Review on microfluidic paper-based analytical devices towards commercialisation. Anal. Chim. Acta 1001, 1–17.

Ballerini, D.R., Li, X., Shen, W., 2012. Patterned paper and alternative materials as substrates for low-cost microfluidic diagnostics. Microfluid. Nanofluidics 13, 769–787. https://doi.org/10.1007/s10404-012-0999-2

Bateman, R.C., Evans, J.A., 1995. Using the glucose oxidase/peroxidase system in enzyme kinetics. J. Chem. Educ. 72, 240–241. https://doi.org/10.1021/ed072pa240

Bhakta, S.A., Borba, R., Taba, M., Garcia, C.D., Carrilho, E., 2014. Determination of nitrite in saliva using microfluidic paper-based analytical devices. Anal. Chim. Acta 809, 117–122. https://doi.org/10.1016/j.aca.2013.11.044

Bjorksten, F., 1970. The Horseradish Peroxidase − Catalyzed Oxidation of Iodide. Outline of the Mechanism. Biochim. Biophys. Acta 212, 396–406.

Brink, S., 1999. Diabetic ketoacidosis. Acta Paediatr. 427, 14–24. https://doi.org/10.1177/1755738017735779

Brooks, T., Keevil, C.W., 1997. A simple artificial urine for the growth of urinary pathogens. Lett. Appl. Microbiol. 24, 203–206. https://doi.org/10.1046/j.1472-765X.1997.00378.x

Brutin, D., Sobac, B., Loquet, B., Sampol, J., 2011. Pattern formation in drying drops of blood. J. Fluid Mech. 667, 85–95. https://doi.org/10.1017/S0022112010005070

Buser, J.R., 2016. Heat, Fluid, and Sample Control in Point-of-Care Diagnostics. ProQuest Diss. Theses. University of Washington.

Cai, J., Yu, B., 2011. A Discussion of the Effect of Tortuosity on the Capillary Imbibition in Porous Media. Transp. Porous Media 89, 251–263. https://doi.org/10.1007/s11242-011-9767-0

Cai, L., Xu, C., Lin, S.H., Luo, J., Wu, M., Yang, F., 2014. A simple paper-based sensor fabricated by selective wet etching of silanized filter paper using a paper mask. Biomicrofluidics 8, 056504(1)–056504(8). https://doi.org/10.1063/1.4898096

Campbell, J.M., Balhoff, J.B., Landwehr, G.M., Rahman, S.M., Vaithiyanathan, M., Melvin, A.T., 2018. Microfluidic and paper-based devices for disease detection and diagnostic research. Int. J. Mol. Sci. 19, 1–38. https://doi.org/10.3390/ijms19092731

Cao, L., Han, G.C., Xiao, H., Chen, Z., Fang, C., 2020. A novel 3D paper-based microfluidic electrochemical glucose biosensor based on rGO-TEPA/PB sensitive film. Anal. Chim. Acta 1096, 34–43. https://doi.org/10.1016/j.aca.2019.10.049

Cate, D.M., Adkins, J.A., Mettakoonpitak, J., Henry, C.S., 2015. Recent Developments in Paper-Based Microfluidic Devices. Anal. Chem. 87, 19–41.

Chaiyo, S., Mehmeti, E., Siangproh, W., Hoang, T.L., Nguyen, H.P., Chailapakul, O., Kalcher, K., 2018. Non-enzymatic electrochemical detection of glucose with a disposable paper-based sensor using a cobalt phthalocyanine–ionic liquid–graphene composite. Biosens. Bioelectron. 102, 113–120. https://doi.org/10.1016/j.bios.2017.11.015

Chang, S., Seo, J., Hong, S., Lee, D.G., Kim, W., 2018. Dynamics of liquid imbibition through paper with intra-fibre pores. J. Fluid Mech. 845, 36–50. https://doi.org/10.1017/jfm.2018.235

Chaudhury, K., Kar, S., Chakraborty, S., 2016. Diffusive dynamics on paper matrix. Appl. Phys. Lett. 109, 224101 (1–5). https://doi.org/10.1063/1.4966992

Chauhan, A., Toley, B.J., 2021. Barrier-Free Microfluidic Paper Analytical Devices for Multiplex Colorimetric Detection of Analytes. Anal. Chem. 93, 8954–8961 https://doi.org/10.1021/acs.analchem.1c01477

Chen, R., Zhang, L., Zang, D., Shen, W., 2016. Blood drop patterns: Formation and applications. Adv. Colloid Interface Sci. 231, 1–14. https://doi.org/10.1016/j.cis.2016.01.008

Chen, X., Chen, J., Wang, F., Xiang, X., Luo, M., Ji, X., He, Z., 2012. Determination of glucose and uric acid with bienzyme colorimetry on microfluidic paper-based analysis devices. Biosens. Bioelectron. 35, 363–368. https://doi.org/10.1016/j.bios.2012.03.018

Chen, Y.-H., Kuo, Z.-K., Cheng, C.-M., 2015. Paper- a potential platform in phamaceutical development. Trends Biotechnol. 33, 4–9.

Chen, Y., Zhang, Y., Liang, Z., Cao, Y., Han, Z., Feng, X., 2020. Flexible inorganic bioelectronics. npj Flex. Electron. 4, 1–20. https://doi.org/10.1038/s41528-020-0065-1

Chin, C.D., Linder, V., Sia, S.K., 2007. Lab-on-a-chip devices for global health: Past studies and future opportunities. Lab Chip 7, 41–57. https://doi.org/10.1039/b611455e

Clayden, J., Greeves, N., Warren, S., 2001. Organic chemistry, Journal of the Chemical Society, Abstracts. https://doi.org/10.1039/ca9161000197

Cummins, B.M., Chinthapatla, R., Ligler, F.S., Walker, G.M., 2017. Time-Dependent Model for Fluid Flow in Porous Materials with Multiple Pore Sizes. Anal. Chem. 89, 4377–4381. https://doi.org/10.1021/acs.analchem.6b04717

Demirel, G., Babur, E., 2014. Vapor-phase deposition of polymers as a simple and versatile technique to generate paper-based microfluidic platforms for bioassay applications. Analyst 139, 2326–2331. https://doi.org/10.1039/c4an00022f

Dolman, D., Newell, G.A., Thurlow, M.D., 1975. A kinetic study of the reaction of horseradish peroxidase with hydrogen peroxide. Can. J. Biochem. 53, 495–501. https://doi.org/10.1139/o75-069

Dou, M., Sanjay, S.T., Benhabib, M., Xu, F., Li, X., 2015. Low-cost bioanalysis on paper-based and its hybrid microfluidic paltforms. Talanta 145, 43–54.

Dunford, H.B., Stillman, J.S., 1976. On the function and mechanism of action of peroxidases. Coord. Chem. Rev. 19, 187–251. https://doi.org/10.1016/S0010-8545(00)80316-1

Dunger, D.B., Sperling, M.A., Acerini, C.L., Bohn, D.J., Daneman, D., Danne, T.P.A., Glaser, N.S., Hanas, R., Hintz, R.L., Levitsky, L.L., Savage, M.O., Tasker, R.C., Wolfsdorf, J.I., 2004. ESPE/LWPES consensus statement on diabetic ketoacidosis in children and adolescents. Arch. Dis. Child. 89, 188–194. https://doi.org/10.1136/adc.2003.044875

Elizalde, E., Urteaga, R., Berli, C.L.A., 2015. Rational design of capillary-driven flows for paper-based microfluidics. Lab Chip 15, 2173–2180. https://doi.org/10.1039/c4lc01487a

Foster, J.R., Morrison, G., Fraser, D.D., 2011. Diabetic ketoacidosis-associated stroke in children and youth. Stroke Res. Treat. 1–12. https://doi.org/10.4061/2011/219706

Fu, L.-M., Wang, Y.-N., 2018. Detection methods and applications of microfluidic paper-based analytical devices. Trends Anal. Chem. 107, 196–211.

Gabriel, E.F.M., Garcia, P.T., Lopes, F.M., Coltro, W.K.T., 2017. Paper-based colorimetric biosensor for tear glucose measurements. Micromachines 8, 1–9. https://doi.org/10.3390/mi8040104

Gong, M.M., Sinton, D., 2017. Turning the Page: Advancing Paper-Based Microfluidics for Broad Diagnostic Application. Chem. Rev. 117, 8447–8480. https://doi.org/10.1021/acs.chemrev.7b00024

Gu, Y., Zhang, T., Chen, H., Wang, F., Pu, Y., Gao, C., Li, S., 2019. Mini Review on Flexible and Wearable Electronics for Monitoring Human Health Information. Nanoscale Res. Lett. 14, 1–15. https://doi.org/10.1186/s11671-019-3084-x

Hawk, 1955. Practical Physiological Chemistry.

Hertaeg, M.J., Tabor, R.F., Berry, J.D., Garnier, G., 2019. Dynamics of stain growth from sessile droplets on paper. J. Colloid Interface Sci. 541, 312–321. https://doi.org/10.1016/j.jcis.2019.01.032

Hu, J., Wang, S.Q., Wang, L., Li, F., Pingguan-Murphy, B., Lu, T.J., Xu, F., 2014. Advances in paper-based point-of-care diagnostics. Biosens. Bioelectron. 54, 585–597. https://doi.org/10.1016/j.bios.2013.10.075

Islam, M.N., Ahmed, I., Anik, M.I., Ferdous, M.S., Khan, M.S., 2018. Developing paper based diagnostic technique to detect uric acid in urine. Front. Chem. 6, 1–12. https://doi.org/10.3389/fchem.2018.00496

JP, C., AJ., G., 1990. Clinical Methods: The History, Physical, and Laboratory Examinations., 3rd editio. ed.

Kaewarsa, P., Laiwattanapaisal, W., Palasuwan, A., Palasuwan, D., 2017. A new paper-based analytical device for detection of Glucose-6-phosphate dehydrogenase deficiency. Talanta 164, 534–539. https://doi.org/10.1016/j.talanta.2016.12.026

Kang, B.H., Park, M., Jeong, K.H., 2017. Colorimetric Schirmer strip for tear glucose detection. Biochip J. 11, 294–299. https://doi.org/10.1007/s13206-017-1405-7

Kap, Ö., Kılıç, V., Hardy, J.G., Horzum, N., 2021. Smartphone-based colorimetric detection systems for glucose monitoring in the diagnosis and management of diabetes. Analyst 146, 2784–2806. https://doi.org/10.1039/d0an02031a

Kar, S., Das, S.S., Laha, S., Chakraborty, S., 2020. Microfluidics on Porous Substrates Mediated by Capillarity-Driven Transport. Ind. Eng. Chem. Res. 59, 3644–3654. https://doi.org/10.1021/acs.iecr.9b04772

Kerl, M.E., 2001. Diabetic Ketoacidosis : Pathophysiology and Clinical and. Compendium 23.

Khan, A.S.A., Talbot, J.A., Tieszen, K.L., Gardener, E.A., Gibson, J.M., New, J.P., 2004. Evaluation of a bedside blood ketone sensor: The effects of acidosis, hyperglycaemia and acetoacetate on sensor performance. Diabet. Med. 21, 782–785. https://doi.org/10.1111/j.1464-5491.2004.01233.x

Khan, S., Ali, S., Bermak, A., 2019. Recent developments in printing flexible and wearable sensing electronics for healthcare applications. Sensors (Switzerland) 19, 1–34. https://doi.org/10.3390/s19051230

Kim, S.J., Kim, D., Kim, S., 2020. Simultaneous quantification of multiple biomarkers on a self-calibrating microfluidic paper-based analytic device. Anal. Chim. Acta 1097, 120–126. https://doi.org/10.1016/j.aca.2019.10.068

Klasner, S.A., Price, A.K., Hoeman, K.W., Wilson, R.S., Bell, K.J., Culbertson, C.T., 2010. Paper-based microfluidic devices for analysis of clinically relevant analytes present in urine and saliva. Anal. Bioanal. Chem. 397, 1821–1829. https://doi.org/10.1007/s00216-010-3718-4

Klocker, A.A., Phelan, H., Twigg, S.M., Craig, M.E., 2013. Blood β-hydroxybutyrate vs. urine acetoacetate testing for the prevention and management of ketoacidosis in Type 1 diabetes: A systematic review. Diabet. Med. 30, 818–824. https://doi.org/10.1111/dme.12136

Kumar, S., Agarwal, A.K., Bhattacharya, S., 2019. Paper Microfluidics Theory and Applications. https://doi.org/10.1007/978-981-15-0489-1_1

Laffel, L., 1999. Ketone bodies: A review of physiology, pathophysiology and application of monitoring to diabetes. Diabetes. Metab. Res. Rev. 15, 412–426. https://doi.org/10.1002/(SICI)1520-7560(199911/12)15:6<412::AID-DMRR72<3.0.CO;2-8

Laios, I., Fast, D.M.., Pardue, H.L., 1986. Development and evaluation of conditions for the kinetic quantitation of acetoacetate in body fluids. Anal. Chim. Acta 180, 429–443.

Laios, I.D., Pardue, H.L., 1993. Kinetic Study of the Reaction of Acetoacetate with Glycine and Sodium Nitroprusside. Anal. Chem. 65, 1903–1909. https://doi.org/10.1021/ac00062a016

Lamas-Ardisana, P.J., Martínez-Paredes, G., Añorga, L., Grande, H.J., 2018. Glucose biosensor based on disposable electrochemical paper-based transducers fully fabricated by screen-printing. Biosens. Bioelectron. 109, 8–12. https://doi.org/10.1016/j.bios.2018.02.061

Leal, L.G., 2007. Advanced Transport Phenomena, Advanced Transport Phenomena. Cambrdige University Press, New York. https://doi.org/10.1017/cbo9780511800238.002

Leitner, C., Volc, J., Haltrich, D., 2001. Purification and Characterization of Pyranose Oxidase from the White Rot Fungus Trametes multicolor. Appl. Environ. Microbiol. 67, 3636–3644. https://doi.org/10.1128/AEM.67.8.3636-3644.2001

Lepowsky, E., Ghaderinezhad, F., Knowlton, S., Tasoglu, S., 2017. Paper - Based Assays for Urine Analysis. Biomicrofluidics 11.

Li, X., Ballerini, D.R., Shen, W., 2012. A perspective on paper-based microfluidics: Current status and future trends. Biomicrofluidics 6, 011301(1)–011301(13). https://doi.org/10.1063/1.3687398

Li, X., Tian, J., Nguyen, T., Shen, W., 2008. Paper-based microfluidic devices by plasma treatment. Anal. Chem. 80, 9131–9134. https://doi.org/10.1021/ac801729t

Lim, W.Y., Goh, B.T., Khor, S.M., 2017. Microfluidoc paper-based analytical devices for potential use in quantitative and direct detection of disease biomarkers in clinical analysis. J. Chromatogr. B 1060, 424–442.

Lin, C., Tseng, C., Chuang, T., Lee, D., Lee, G., S, C.L.B., 2011. Urine analysis in microfluidic devices 2669–2688. https://doi.org/10.1039/c1an15029d

Lin, C.C., Tseng, C.C., Chuang, T.K., Lee, D.S., Lee, G. Bin, 2011. Urine analysis in microfluidic devices. Analyst 136, 2669–2688. https://doi.org/10.1039/c1an15029d

Liu, M.M., Lian, X., Liu, H., Guo, Z.Z., Huang, H.H., Lei, Y., Peng, H.P., Chen, W., Lin, X.H., Liu, A.L., Xia, X.H., 2019. A colorimetric assay for sensitive detection of hydrogen peroxide and glucose in microfluidic paper-based analytical devices integrated with starch-iodide-gelatin system. Talanta 200, 511–517. https://doi.org/10.1016/j.talanta.2019.03.089

Liu, Z., Hu, J., Zhao, Y., Qu, Z., Xu, F., 2015. Experimental and numerical studies on liquid wicking into filter papers for paper-based diagnostics. Appl. Therm. Eng. 88, 280–287. https://doi.org/10.1016/j.applthermaleng.2014.09.057

Lucas, R., 1918. Ueber das Zeitgesetz des kapillaren Aufstiegs von Flüssigkeiten. Kolloid-Zeitschrift 23, 15–22. https://doi.org/10.1007/BF01461107

MacDonald, B.D., 2018. Flow of liquids through paper. J. Fluid Mech. 852, 1–4. https://doi.org/10.1017/jfm.2018.536

Mahato, K., Srivastava, A., Chandra, P., 2017. Paper based diagnostics for personalised health care: Emerging technologies and commercial aspects. Biosens. Bioelectron. 8, 246–259.

Mao, X., Huang, T.J., 2012. Microfluidic diagnostics for the developing world. Lab Chip 12, 1412–1416. https://doi.org/10.1039/c2lc90022j

Martinez, A.W., Phillips, S.T., Butte, M.J., Whitesides, G.M., 2007. Patterned paper as a platform for inexpensive, low-volume, portable bioassays. Angew. Chemie - Int. Ed. 46, 1318–1320. https://doi.org/10.1002/anie.200603817

Martinez, A.W., Phillips, S.T., Carrilho, E., Thomas, S.W., Sindi, H., Whitesides, G.M., 2008a. Simple telemedicine for developing regions: Camera phones and paper-based microfluidic devices for real-time, off-site diagnosis. Anal. Chem. 80, 3699–3707. https://doi.org/10.1021/ac800112r

Martinez, A.W., Phillips, S.T., Nie, Z., Cheng, C.M., Carrilho, E., Wiley, B.J., Whitesides, G.M., 2010a. Programmable diagnostic devices made from paper and tape. Lab Chip 10, 2499–2504. https://doi.org/10.1039/c0lc00021c

Martinez, A.W., Phillips, S.T., Whitesides, G.M., 2008b. Three-dimensional microfluidic devices fabricated in layered paper and tape. Proc. Natl. Acad. Sci. U. S. A. 105, 19606–19611. https://doi.org/10.1073/pnas.0810903105

Martinez, A.W., Phillips, S.T., Whitesides, G.M., Carrilho, E., 2010b. Diagnostics for the developing world: Microfluidic paper-based analytical devices. Anal. Chem. 82, 3–10. https://doi.org/10.1021/ac9013989

Millington, R.J., 1959. Gas Diffusion in Porous Media. Science (80-.). 130, 100–102.

Millington, R.J., Quirk, J.P., 1959. Permeability of Porous Media. Nature 183, 387–388.

Modha, S., Castro, C., Tsutsui, H., 2021. Recent developments in flow modeling and fluid control for paper-based microfluidic biosensors. Biosens. Bioelectron. 178, 113026(1–17). https://doi.org/10.1016/j.bios.2021.113026

Molina, F., Valencia, M.C., De La Torre, J., Capitan-Vallvey, F., 1989. Determination of acetoacetate in urine by solid-phase spectrophotometry. J. Pharm. Biomed. Anal. 7, 843–850.

Mukhopadhyay, M., Ray, R., Ayushman, M., Sood, P., Bhattacharyya, M., Sarkar, D., DasGupta, S., 2020. Interfacial energy driven distinctive pattern formation during the drying of blood droplets. J. Colloid Interface Sci. 573, 307–316. https://doi.org/10.1016/j.jcis.2020.04.008

Musile, G., Agard, Y., Wang, L., Palo, E.F., McCord, B., Tagliaro, F., 2021. Paper-based microfluidic devices: On-site tools for crime scene investigation. Trends Anal. Chem. 143, 116406(1)–116406(15).

Nery;, E.W., Kubota, L.T., 2016. Evaluation of enzyme immobilization methods for paper-based devices—A glucose oxidase study.pdf. J. Pharm. Biomed. Anal. 117, 551–559.

Ngo, Y.L.T., Nguyen, P.L., Jana, J., Choi, W.M., Chung, J.S., Hur, S.H., 2021. Simple paper-based colorimetric and fluorescent glucose sensor using N-doped carbon dots and metal oxide hybrid structures. Anal. Chim. Acta 1147, 187–198. https://doi.org/10.1016/j.aca.2020.11.023

Nie, Z., Nijhuis, C.A., Gong, J., Chen, X., Kumachev, A., Martinez, A.W., Narovlyansky, M., Whitesides, G.M., 2010. Electrochemical sensing in paper-based microfluidic devices. Lab Chip 10, 477–483. https://doi.org/10.1039/b917150a

Nishat, S., Jafry, A.T., Martinez, A.W., Awan, F.R., 2021. Paper-based microfluidics: Simplified fabrication and assay methods. Sensors Actuators, B Chem. 336, 129681. https://doi.org/10.1016/j.snb.2021.129681

Olkkonen, J., Lehtinen, K., Erho, T., 2010. Flexographically printed fluidic structures in paper. Anal. Chem. 82, 10246–10250. https://doi.org/10.1021/ac1027066

Ozaki, S. ichi, de Montellano, P.R.O., 1995. Molecular Engineering of Horseradish Peroxidase: Thioether Sulfoxidation and Styrene Epoxidation by Phe-41 Leucine and Threonine Mutants. J. Am. Chem. Soc. 117, 7056–7064. https://doi.org/10.1021/ja00132a003

Park, H.D., Lee, K.J., Yoon, H.R., Nam, H.H., 2005. Design of a portable urine glucose monitoring system for health care. Comput. Biol. Med. 35, 275–286. https://doi.org/10.1016/j.compbiomed.2004.02.003

Parsa, M., Harmand, S., Sefiane, K., 2018. Mechanisms of pattern formation from dried sessile drops. Adv. Colloid Interface Sci. 254, 22–47. https://doi.org/10.1016/j.cis.2018.03.007

Patari, S., Mahapatra, P.S., 2020. Liquid wicking in a paper strip: An experimental and numerical study. ACS Omega 5, 22931–22939. https://doi.org/10.1021/acsomega.0c02407

Pedersen, K.J., 1961. The Hydrolysis of Ethyl Acetoacetate and the Decarboxylation of Acetoacetic Acid in Strongly Acid Solution. Acta Chem. Scand. 15, 1718–1722.

Rath, D., Sathishkumar, N., Toley, B.J., 2018. Experimental Measurement of Parameters Governing Flow Rates and Partial Saturation in Paper-Based Microfluidic Devices. Langmuir 34, 8758–8766. https://doi.org/10.1021/acs.langmuir.8b01345

Rath, D., Toley, B.J., 2021. Modeling-Guided Design of Paper Microfluidic Networks: A Case Study of Sequential Fluid Delivery. ACS Sensors 6, 91–99. https://doi.org/10.1021/acssensors.0c01840

Richards, L.A., 1931. Capillary conduction of liquids through porous mediums. Physics (College. Park. Md). 1, 318–333. https://doi.org/10.1063/1.1745010

Roman;, R., Dunford, H.B., 1972. pH Dependence of the Oxidation of Iodide by Compound I of Horseradish Peroxidase. Biochemistry 11, 2076–2082.

Rossini;, E.L., Milani;, M.I., Lima;, L.S., Pezza, H.R., 2021. Paper microfluidic device using carbon dots to detect glucose and lactate in saliva samples. Spectrochim. Acta part A Mol. Biomol. Spectrosc. 248, 119285 (1–7).

Rossini, E.L., Milani, M.I., Pezza, H.R., 2019. Green synthesis of fluorescent carbon dots for determination of glucose in biofluids using a paper platform. Talanta 201, 503–510. https://doi.org/10.1016/j.talanta.2019.04.045

Sechi, D., Greer, B., Johnson, J., Hashemi, N., 2013. Three-dimensional paper-based microfluidic device for assays of protein and glucose in urine. Anal. Chem. 85, 10733–10737. https://doi.org/10.1021/ac4014868

Sefiane, K., 2010. On the formation of regular patterns from drying droplets and their potential use for bio-medical applications. J. Bionic Eng. 7, S82–S93. https://doi.org/10.1016/S1672-6529(09)60221-3

Selvakumar, B., Kathiravan, A., 2021. Sensory materials for microfluidic paper based analytical devices - A review. Talanta 235, 1–15.

Sharp, V.J.A., Antes, L.M., Sanders, M.L., Lockwood, G.M., 2008. Urine Tests, Handbook of Veterinary Nursing. https://doi.org/10.1002/9780470690376.ch35

Shen, L., Hagen, J.A., Papautsky, I., 2012. Point-of-care colorimetric detection with a smartphone. Lab Chip 12, 4240–4243. https://doi.org/10.1039/c2lc40741h

Shou, D., Ye, L., Fan, J., Fu, K., Mei, M., Wang, H., Chen, Q., 2014. Geometry-induced asymmetric capillary flow. Langmuir 30, 5448–5454. https://doi.org/10.1021/la500479e

Singh, M., Ungku Faiz, U.Z.A., Gravelsins, S., Suganuma, Y., Kotoulas, N.K., Croxall, M., Khan-Trottier, A., Goh, C., Dhirani, A.-A., 2021. Glucose oxidase kinetics using MnO 2 nanosheets: confirming Michaelis–Menten kinetics and quantifying decreasing enzyme performance with increasing buffer concentration. Nanoscale Adv. 3, 3816–3823. https://doi.org/10.1039/d1na00311a

Soni, A., Jha, S.K., 2015. A paper strip based non-invasive glucose biosensor for salivary analysis. Biosens. Bioelectron. 67, 763–768. https://doi.org/10.1016/j.bios.2014.09.042

Starov, V.M., Kostvintsev, S.R., Sobolev, V.D., Velarde, M.G., Zhdanov, S.A., 2002. Spreading of Liquid Drops over Dry Porous Layers : Complete Wetting Case. J. Colloid Interface Sci. 252, 397–408. https://doi.org/10.1006/jcis.2002.8450

Svendsen, W.E., 2015. Lab-on-a-Chip Devices and Micro-Total Analysis Systems, Springer.

Tai, W., Chang, Y., Chou, D., Fu, L., 2021. Lab-on-Paper Devices for Diagnosis of Human Diseases Using Urine Samples - A Review. Biosensors 11, 1–23.

Tam, T. Van, Hur, S.H., Chung, J.S., Choi, W.M., 2021. Novel paper- and fiber optic-based fluorescent sensor for glucose detection using aniline-functionalized graphene quantum dots. Sensors Actuators, B Chem. 329, 129250. https://doi.org/10.1016/j.snb.2020.129250

Tao, Z., Raffel, R.A., Souid, A.K., Goodisman, J., 2009. Kinetic studies on enzyme-catalyzed reactions: Oxidation of glucose, decomposition of hydrogen peroxide and their combination. Biophys. J. 96, 2977–2988. https://doi.org/10.1016/j.bpj.2008.11.071

Tseng, C.C., Kung, C. Te, Chen, R.F., Tsai, M.H., Chao, H.R., Wang, Y.N., Fu, L.M., 2021. Recent advances in microfluidic paper-based assay devices for diagnosis of human diseases using saliva, tears and sweat samples. Sensors Actuators, B Chem. 342, 130078. https://doi.org/10.1016/j.snb.2021.130078

Ugarova;, N.N., Lebedeva;, O. V., Berezin, L. V., 1981. Horseradish Peroxidase Catalysis I Steady-state Kinetics of Peroxidase-Catalyzed Individual and Co-oxidation of Potassium Ferrocyanide and O-Dianisidine by Hydrogen Peroxide. J. Mol. Catal. 13, 215–225.

Verma, A., Sharma, A., 2010. Enhanced Self-Organized Dewetting of Ultrathin Polymer Films Under Water-Organic Solutions: Fabrication of Sub-micrometer Spherical Lens Arrays. Adv. Mater 22, 5306–5309. https://doi.org/10.1002/adma.201002768

Wallace, T.M., Matthews, D.R., 2004. Recent advances in the monitoring and management of diabetic ketoacidosis. QJM - Mon. J. Assoc. Physicians 97, 773–780. https://doi.org/10.1093/qjmed/hch132

Wang, C.C., Hennek, J.W., Ainla, A., Kumar, A.A., Lan, W.J., Im, J., Smith, B.S., Zhao, M., Whitesides, G.M., 2016. A Paper-Based Pop-up Electrochemical Device for Analysis of Beta-Hydroxybutyrate. Anal. Chem. 88, 6326–6333. https://doi.org/10.1021/acs.analchem.6b00568

Washburn, E.D., 1921. The Dynamics of Capillary Flow. Phys. Rev. 17, 273–283. https://doi.org/10.1103/PhysRev.18.206

Whitesides, G.M., 2006. The origins and the future of microfluidics. Nature 442, 368–373. https://doi.org/10.1038/nature05058

WHO Global Report on Diabetes, 2016. Global Report on Diabetes. Isbn 978, 6–86.

William, D.M., 1998. Analysis of Transport Phenomena, Oxford University Press. Oxford University Press, New York.

Wolfsdorf, J., Glaser, N., Sperling, M.A., 2006. Diabetic ketoacidosis in infants, children, and adolescents: A consensus statement from the American Diabetes Association. Diabetes Care 29, 1150–1159. https://doi.org/10.2337/dc06-9909

Xia, Y., Si, J., Li, Z., 2016. Fabrication techniques for microfluidic paper-based analytical devices and their applications for biological testing: A review. Biosens. Bioelectron. 77, 774–789. https://doi.org/10.1016/j.bios.2015.10.032

Yamada, K., Shibata, H., Suzuki, K., Citterio, D., 2017. Toward practical application of paper-based microfluidics for medical diagnostics: state-of-the-art and challenges. Lab Chip 17, 1206–1249. https://doi.org/10.1039/c6lc01577h

Yetisen, A.K., Akram, M.S., Lowe, C.R., 2013. Paper-based microfluidic point-of-care diagnostic devices. Lab Chip 13, 2210–2251. https://doi.org/10.1039/c3lc50169h

Yetisen, A.K., Jiang, N., Tamayol, A., Ruiz-Esparza, G.U., Zhang, Y.S., Medina-Pando, S., Gupta, A., Wolffsohn, J.S., Butt, H., Khademhosseini, A., Yun, S.H., 2017. Paper-based microfluidic system for tear electrolyte analysis. Lab Chip 17, 1137–1148. https://doi.org/10.1039/c6lc01450j

Zhang, H., Chen, Z., Dai, J., Zhang, W., Jiang, Y., Zhou, A., 2021. A low-cost mobile platform for whole blood glucose monitoring using colorimetric method. Microchem. J. 162, 105814(1)–105814(7). https://doi.org/10.1016/j.microc.2020.105814

Zhang, H., Smith, E., Zhang, W., Zhou, A., 2019. Inkjet printed microfluidic paper-based analytical device (μPAD) for glucose colorimetric detection in artificial urine. Biomed. Microdevices 21, 1–10. https://doi.org/10.1007/s10544-019-0388-7

Zhu, W.J., Feng, D.Q., Chen, M., Chen, Z.D., Zhu, R., Fang, H.L., Wang, W., 2014. Bienzyme colorimetric detection of glucose with self-calibration based on tree-shaped paper strip. Sensors Actuators, B Chem. 190, 414–418. https://doi.org/10.1016/j.snb.2013.09.007

