## Supplementary Section for the Manuscript for "Design, Fabrication, and Theoretical Investigation of a Cost-Effective Laser Printing Based Colorimetric Paper Sensor for Non-Invasive Glucose and Ketone Detection"

### Supplemental Section

*[#]Equal contribution*

*[*]Corresponding author*

Ph.: +91 - 3222 – 283922

### Description of the Supplemental Sections:

S1: Characteristics of the filter papers

S2: Optimization of the Device Design for Glucose and Ketone Sensors

S3: Schematic and parameters for the COMSOL simulation

S4: Parameter estimation

S5: Comparison between the image intensity ratios between ImageJ and RGAD app.

### S1. Characteristics of the filter papers:

**Table S1:** Comparison of filter papers tested in the present work (Kumar et al., 2019)

| Substrate | Cost(₹)(110mm diameter circles) | Thickness (mm) |
|---|---|---|
| Whatman filter paper Grade 113 | 17.9 per piece | 0.42 |
| Whatman filter paper Grade 43 | 30.80 per piece | 0.22 |
| Whatman filter paper Grade 4 | 9.9 per piece | 0.21 |
| Whatman filter paper Grade 3 | 12.9 per piece | 0.39 |
| Whatman filter paper Grade 1 | 7.9 per piece | 0.18 |

### S2. Optimization of the Device Design for Glucose and Ketone Sensors:

Design 1 was a simple circular region surrounded by the hydrophobic barriers for spotting the reagents and the analyte. In this case, low contents of glucose resulted in the development of a very faint colour at the periphery which was difficult to capture, even under a microscope. In order to intensify the colour, sharp edges were incorporated in design 2. The results were similar to that observed previously. Understanding that the reaction requires a channel so that the produced low molecular weight iodine molecules could be deposited as the flow proceeds, a channel was incorporated in the next design (design 3). The results were significantly improved but the colour was deposited only at the periphery. With the inclusion of sharp edges in design 4, pixilation of the edges due to the printing rendered this design unfavourable for the assay. Hence, the number of sharp edges were subsequently increased which finally led to design 7 with a separate zone for detection. This design showed even distribution of colour in the detection area, which could easily be captured. Thus, this was chosen as final design for the glucose assay.

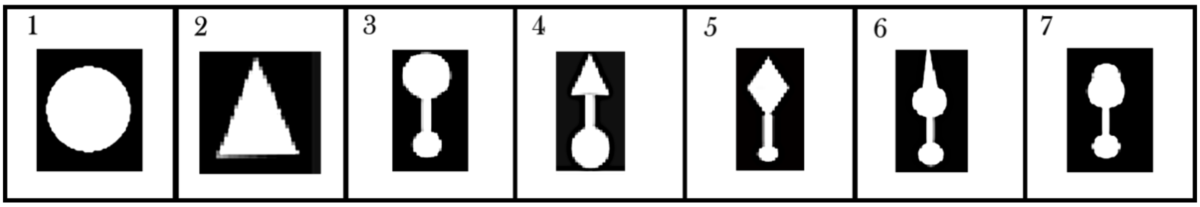


Figure S1: Progression of device designs considered for the present work

### S3. Schematic and Parameters for the COMSOL Simulation:

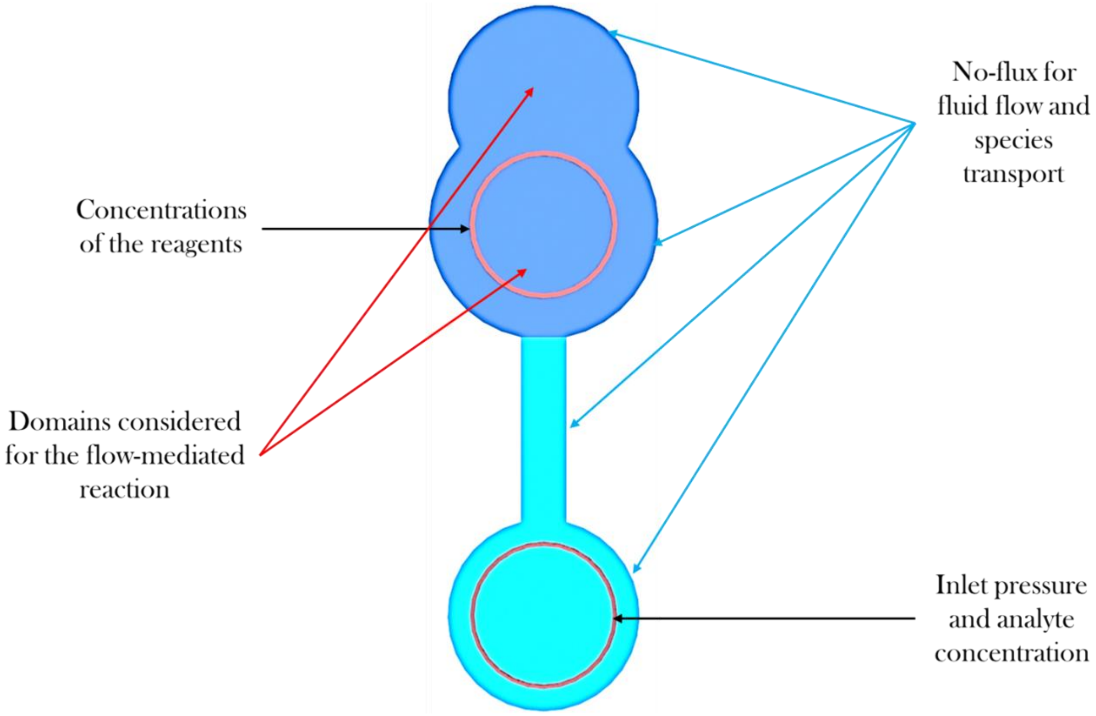


Figure S2: Schematic depicting the reactions domains, the inlet and boundary conditions chosen for simulating the glucose sensor.

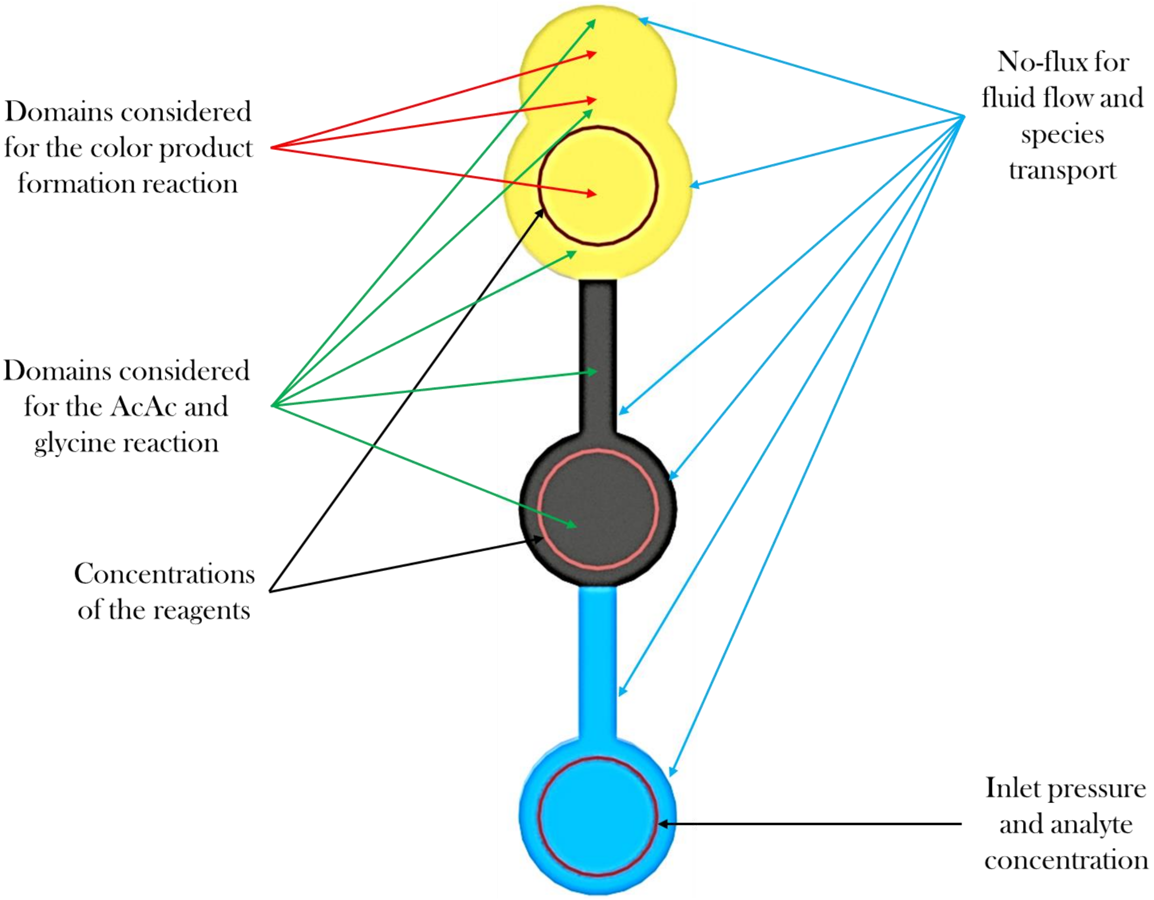


Figure S3: Schematic depicting the inlet and boundary conditions, along with the selected domains for the two reactions pertaining to the simulating of the AcAc sensor.

Table S2: Parameters used for the COMSOL simulation

| Parameter | Value | Description | Reference |
|---|---|---|---|
| Epsln | 0.7 | Porosity of the filter paper | (Chaudhury et al., 2016; Chauhan and Toley, 2021; Kar et al., 2020) |
| Alpha | 1 | Van Genutchen parameter_1 | (Buser, 2016; Rath et al., 2018) |

| | | | |
|---|---|---|---|
| q | 1.30 | Van Genutchen parameter_2 | (Buser, 2016; Rath et al., 2018) |
| l | 0.03 | Van Genutchen parameter_3 | (Buser, 2016; Rath et al., 2018) |
| thckns | 0.18 [mm] | Thickness of the filter paper | Table S1 |
| rho_liq | 996.57 kg/m^3] | Density of the liquid | - |
| mu_liq | 8.502E-4 [Pa. s] | Dynamic viscosity of the liquid | - |
| gamma | 0.0723 [N/m] | Surface tension of the liquid | - |
| theta_r | 1E-4 | Residual saturation | Assumption giving due consideration to the bound water content in the paper matrix (Buser, 2016) |
| kappa_s | 1.87E-6 [m/s] | Hydraulic conductivity at 100 % saturation | (Buser, 2016; Rath et al., 2018) |
| T_isothrml | 300.15 [K] | System temperature | Experimental condition |
| P0 | -101.325 [kPa] | Initial pressure throughout the filter paper matrix | Experimental condition and based on (Buser, 2016; Rath et al., 2018; |

| | | | |
|---|---|---|---|
| | | | Rath and Toley, 2021) |

### S4. Parameter estimation:

Table S3: Estimation of parameters for 99% confidence interval for the glucose assay

| S. No. | Concentration range (in mM) | $\overline{x}$ | $\sigma$ | $n$ | Lower Limit | Upper Limit |
|---|---|---|---|---|---|---|
| 1. | 1 | 0.634 | 0.023 | 30 | 0.623 | 0.645 |
| 2. | 2 | 0.593 | 0.015 | 30 | 0.586 | 0.599 |
| 3. | 3 | 0.575 | 0.004 | 30 | 0.573 | 0.577 |
| 4. | 4 | 0.566 | 0.003 | 30 | 0.564 | 0.567 |
| 5. | 5 | 0.554 | 0.004 | 30 | 0.552 | 0.556 |
| 6. | 6 | 0.545 | 0.006 | 30 | 0.542 | 0.547 |
| 7. | 7 | 0.536 | 0.005 | 30 | 0.534 | 0.539 |
| 8. | 8 | 0.524 | 0.004 | 30 | 0.522 | 0.526 |
| 9. | 9 | 0.511 | 0.005 | 30 | 0.509 | 0.513 |
| 10. | 10 | 0.493 | 0.003 | 30 | 0.491 | 0.494 |
| 11. | 11 | 0.478 | 0.006 | 30 | 0.475 | 0.480 |
| 12. | 12 | 0.468 | 0.004 | 30 | 0.466 | 0.470 |
| 13. | 13 | 0.453 | 0.004 | 30 | 0.451 | 0.454 |
| 14. | 14 | 0.435 | 0.006 | 30 | 0.433 | 0.438 |
| 15. | 15 | 0.414 | 0.018 | 30 | 0.405 | 0.422 |

Table S4: Estimation of parameters for 99% confidence interval for the ketone assay

| S. No. | Concentration (in mM) | $\bar{x}$ | $\sigma$ | $n$ | Lower Limit | Upper Limit |
|---|---|---|---|---|---|---|
| 1. | 1 | 0.723 | 0.024 | 30 | 0.711 | 0.734 |
| 2. | 2 | 0.657 | 0.031 | 30 | 0.642 | 0.672 |
| 3. | 3 | 0.578 | 0.028 | 30 | 0.565 | 0.591 |
| 4. | 4 | 0.545 | 0.029 | 30 | 0.532 | 0.559 |
| 5. | 5 | 0.522 | 0.025 | 30 | 0.510 | 0.533 |
| 6. | 6 | 0.503 | 0.022 | 30 | 0.493 | 0.514 |
| 7. | 7 | 0.492 | 0.034 | 30 | 0.479 | 0.504 |
| 8. | 8 | 0.481 | 0.027 | 30 | 0.465 | 0.497 |
| 9. | 9 | 0.431 | 0.027 | 30 | 0.418 | 0.444 |
| 10. | 10 | 0.435 | 0.041 | 30 | 0.416 | 0.454 |

### S5. Comparison between the image intensity ratios between ImageJ and RGAD app.:

Table S5: Comparison for glucose assay

| S. No | Intensity ratio obtained from the MATLAB app. (Glucose) | Intensity ratio obtained from ImageJ (Glucose) | Category based on Glucose Content |
|---|---|---|---|
| 1 | 0.563 | 0.564 | Moderate |
| 2 | 0.477 | 0.473 | Very High |

| | | | |
|---|---|---|---|
| 3 | 0.637 | 0.638 | Normal |
| 4 | 0.411 | 0.412 | Alarming |
| 5 | 0.550 | 0.550 | Moderate |
| 6 | 0.491 | 0.493 | Very High |
| 7 | 0.664 | 0.666 | Normal |
| 8 | 0.451 | 0.452 | Very High |
| 9 | 0.574 | 0.574 | Moderate |
| 10 | 0.432 | 0.433 | Alarming |

Table S6: Comparison for AcAc assay

| S. No | Intensity ratio obtained from the MATLAB app. (Ketone) | Intensity ratio obtained from ImageJ (Ketone) | Category based on Ketone Content |
|---|---|---|---|
| 1 | 0.513 | 0.515 | High |
| 2 | 0.721 | 0.719 | Normal |
| 3 | 0.431 | 0.430 | Very High |
| 4 | 0.509 | 0.511 | High |
| 5 | 0.728 | 0.731 | Normal |
| 6 | 0.421 | 0.422 | Very High |
| 7 | 0.542 | 0.541 | Moderate |
| 8 | 0.636 | 0.639 | Moderate |
| 9 | 0.46 | 0.462 | Very High |
| 10 | 0.581 | 0.583 | Moderate |

### References:


Buser, J.R., 2016. Heat, Fluid, and Sample Control in Point-of-Care Diagnostics. ProQuest Diss. Theses. University of Washington.

Chaudhury, K., Kar, S., Chakraborty, S., 2016. Diffusive dynamics on paper matrix. Appl. Phys. Lett. 109, 224101 (1–5). https://doi.org/10.1063/1.4966992

Chauhan, A., Toley, B.J., 2021. Barrier-Free Microfluidic Paper Analytical Devices for Multiplex Colorimetric Detection of Analytes. Anal. Chem. 93, 8954–8961. https://doi.org/10.1021/acs.analchem.1c01477

Kar, S., Das, S.S., Laha, S., Chakraborty, S., 2020. Microfluidics on Porous Substrates Mediated by Capillarity-Driven Transport. Ind. Eng. Chem. Res. 59, 3644–3654. https://doi.org/10.1021/acs.iecr.9b04772

Kumar, S., Agarwal, A.K., Bhattacharya, S., 2019. Paper Microfluidics Theory and Applications. https://doi.org/10.1007/978-981-15-0489-1_1

Rath, D., Sathishkumar, N., Toley, B.J., 2018. Experimental Measurement of Parameters Governing Flow Rates and Partial Saturation in Paper-Based Microfluidic Devices. Langmuir 34, 8758–8766. https://doi.org/10.1021/acs.langmuir.8b01345

Rath, D., Toley, B.J., 2021. Modeling-Guided Design of Paper Microfluidic Networks: A Case Study of Sequential Fluid Delivery. ACS Sensors 6, 91–99. https://doi.org/10.1021/acssensors.0c01840
